# Reliability of the TMS-evoked potential in dorsolateral prefrontal cortex

**DOI:** 10.1101/2023.09.04.556283

**Authors:** Juha Gogulski, Christopher C. Cline, Jessica M. Ross, Sara Parmigiani, Corey J. Keller

**Author notes:** Correspondence: Corey Keller, MD, PhD Stanford University, Department of Psychiatry and Behavioral Sciences, 401 Quarry Road, Stanford, CA 94305-5797.

## Abstract

**Background:** We currently lack a robust and reliable method to probe cortical excitability noninvasively from the human dorsolateral prefrontal cortex (dlPFC), a region heavily implicated in psychiatric disorders. We recently found that the strength of *early* and *local* dlPFC single pulse transcranial magnetic stimulation (TMS)-evoked potentials (EL-TEPs) varied widely depending on the anatomical subregion probed, with more medial regions eliciting stronger responses than anterolateral sites. Despite these differences in *amplitude* of response, the *reliability* at each target is not known.

**Objective:** To evaluate the reliability of EL-TEPs across the dlPFC.

**Methods:** In 15 healthy subjects, we quantified within-session reliability of dlPFC EL-TEPs after single pulse TMS to six dlPFC subregions. We evaluated the concordance correlation coefficient (CCC) across targets and analytical parameters including time window, quantification method, region of interest, sensor-vs. source-space, and number of trials.

**Results:** At least one target in the anterior and posterior dlPFC produced reliable EL-TEPs (CCC>0.7). The medial target was most reliable (CCC = 0.78) and the most anterior target was least reliable (CCC = 0.24). ROI size and type (sensor vs. source space) did not affect reliability. Longer (20-60 ms, CCC = 0.62) and later (30-60 ms, CCC = 0.61) time windows resulted in higher reliability compared to earlier and shorter (20-40 ms, CCC 0.43; 20-50 ms, CCC = 0.55) time windows. Peak-to-peak quantification resulted in higher reliability than the mean of the absolute amplitude. Reliable EL-TEPs (CCC up to 0.86) were observed using only 25 TMS trials for a medial dlPFC target.

**Conclusions:** Medial TMS location, wider time window (20-60ms), and peak-to-peak quantification improved reliability. Highly reliable EL-TEPs can be extracted from dlPFC after only a small number of trials.

**Highlights:**

- Medial dlPFC target improved EL-TEP reliability compared to anterior targets.
- After optimizing analytical parameters, at least one anterior and one posterior target was reliable (CCC>0.7).
- Longer (20-60 ms) and later (30-60 ms) time windows were more reliable than earlier and shorter (20-40 ms or 20-50 ms) latencies.
- Peak-to-peak quantification resulted in higher reliability compared to the mean of the absolute amplitude.
- As low as 25 trials can yield reliable EL-TEPs from the dlPFC.

## 1. Introduction

Although transcranial magnetic stimulation (TMS) to the dorsolateral prefrontal cortex (dlPFC) is an effective treatment for depression (1), the neural response to TMS is not fully understood. Developing a marker of cortical excitability measurable in the clinic that can 1) stratify patients into different treatment protocols and 2) adapt treatment in real-time using closed-loop methods, would represent significant advances. One such promising metric is derived from TMS coupled with electroencephalography (TMS-EEG), which produces a causal measure of cortical excitability (2,3). Here, the ‘*EL-TEP’* or the *local* and short-latency (*early,* 20-60 ms) response to single TMS pulses applied to the dlPFC (4) represents an exciting interpretable neurophysiological metric with neural underpinnings (5). EL-TEPs may reflect aspects of prefrontal GABAergic inhibition and NMDA receptor function (6), but these studies need to be further validated and tested in the dlPFC. Furthermore, pilot studies suggest dlPFC EL-TEPs may track with depression pathophysiology (7) and clinical response to TMS (8,9). In summary, EL-TEPs represent an exciting potential brain biomarker for TMS. As multiple dlPFC subregions represent potential novel treatment targets (10–13), understanding how the EL-TEP changes across dlPFC subregions is critical prior to use as a clinical biomarker.

Recent work in our lab investigated the effect of applying single TMS pulses at different dlPFC subregions on EL-TEP amplitude (4). We hypothesized that because the anterolateral portion of the dlPFC activates scalp muscle that induces large early artifacts in EEG (0-20 ms after a single pulse of TMS (14)) which are difficult to remove, the ‘true’ underlying neural response (e.g. EL-TEPs) may be unintentionally masked or suppressed by offline removal techniques and therefore these targets might produce smaller recorded EL-TEPs. Relatedly, we hypothesized that larger EL-TEPs would be observed from stimulation of the posterior and medial dlPFC due to weaker muscle activation (and therefore smaller masking artifacts). Our findings confirmed these hypotheses, with posterior and medial targets eliciting larger EL-TEPs compared to anterolateral targets (4). However, stronger EL-TEP amplitudes at specific subregions importantly do not necessarily translate to higher reliability. To date, the reliability of EL-TEPs across dlPFC subregions has not been characterized but is critically important to study for prefrontal physiology, pathophysiology, and neuromodulation (15). Furthermore, the effect of analytical parameters including analysis time window and region of interest, how the evoked potential is quantified (e.g. peak-to-peak or mean of the absolute amplitude), and the minimal number of trials needed to obtain a reliable prefrontal EL-TEP, have to date not been characterized. While previous studies have evaluated the reliability of prefrontal TEPs (16,17), this question is worth re-investigating due to an improved understanding of the neural and non-neural artifacts that are present in the TEP. Later (>100 ms) prefrontal TEPs are consistently reliable (16–19), but due to the now well-known contribution of off-target sensory activation (20,21), much of this reliability may be of non-specific sensory brain responses to both the auditory click and tactile sensation to TMS. Reliability measures from earlier components of the TEP (e.g. EL-TEP), less likely to be sensory in nature, have been mixed (16,17). Kerwin *et al*. (17) reported moderate reliability (CCC < 0.6) of early TEPs (the N40 peak) within the same day across different blocks. Conversely, Lioumis *et al*. (16) observed strong correlations for early dlPFC TEP peaks when comparing assessments across different days. For these reasons, and because neither study thoroughly investigated the effects of dlPFC subregion and analytic parameters on reliability, we chose to revisit these critical questions.

In this study, we sought to characterize reliability of EL-TEPs across clinically relevant dlPFC stimulation targets and analytic parameters including ROI size, type (sensor and source space), time window, and amplitude quantification method. Additionally, we evaluated the minimum number of trials needed to obtain highly reliable prefrontal EL-TEPs (CCC>0.7). We focused on within-day reliability to provide reliability thresholds for studies that quantify the acute single-day neural effects of TMS. Here, we hypothesized that 1) because EL-TEP reliability may scale with amplitude of response, a TMS target in posterior and medial aspect of the dlPFC would yield the most reliable prefrontal EL-TEPs (4); 2) how the EL-TEP is quantified (*analytic parameters*) would greatly impact reliability; 3) at least 100 trials would be required for a highly reliable prefrontal EL-TEP (17). Our results demonstrated that 1) the most medial dlPFC target was as predicted the most reliable, while the most anterior target was the least reliable; 2) ROI size did not significantly impact reliability; 3) analytic parameters affected reliability, with later (e.g. starting 30 ms after TMS) and longer (e.g. 40 ms) time windows, peak-to-peak quantification method, and sensor-space yielding more reliable EL-TEPs; and 4) TMS to the most medial dlPFC subregion delivered reliable EL-TEPs with as few as 25 single pulses. These findings demonstrate that the EL-TEP can be reliably measured, and that the choice of stimulation target and analytical parameters can greatly impact EL-TEP reliability.

## 2. Methods

### 2.1. Participants

22 healthy participants (23-64 years old, mean=39.45, SD=13.00, 7F/15M) were recruited into the study and provided written informed consent under a protocol approved by the Stanford University Institutional Review Board. Details on the demographics have been reported previously (4). Briefly, the participants were aged 18-65, fluent in English, fully vaccinated against COVID-19, and without moderate or severe depression. Exclusion criteria included psychiatric/neurological disorders including moderate or severe depression: QIDS ≥11 (22), substance abuse during the month before screening, cardiac events, current pregnancy, contraindications for TMS or MRI (23,24), and lifetime history of psychotropic medication use. Out of the initial participants, 6 were excluded (3 before the experimental session, 1 for TMS intolerability assessed by delivering test single pulses before the experiment, 1 for lower back pain, and 2 for technical difficulties), leaving 15 subjects for analysis after completing MRI and TMS-EEG sessions.

### 2.2. Transcranial Magnetic Stimulation

Single-pulse TMS was delivered using a MagVenture Cool-B65 A/P figure-of-eight coil with a MagPro X100 stimulator (MagVenture, Denmark). A thin (0.5 mm) foam pad was attached to the TMS coil to minimize electrode movement, scalp sensation, and bone-conducted auditory response (25). The TMS coil was held and positioned automatically by a robotic arm (TMS Cobot, Axilum Robotics, France). MRI-based neuronavigation (Localite TMS Navigator, Localite, Germany) was used to align the coil to pre-planned individualized coil orientations for each stimulation target (see section 2.2.1). To estimate resting motor threshold (rMT), biphasic single pulses of TMS were delivered to the hand region of the left motor cortex with the coil held tangentially to the scalp and at 45° from midline (26,27). The optimal motor hotspot was defined as the coil position from which TMS produced the largest and most consistent visible twitch in relaxed right first dorsal interosseous muscle (FDI; (27)). rMT was determined to be the minimum intensity that elicited a visible twitch in relaxed FDI in ≥ 5 out of 10 pulses (28,29). Stimulation intensity was adjusted relative to motor threshold based on electric field strength at the cortical surface, as reported previously (4).

#### 2.2.1 dlPFC Target Selection

For each subject, single pulses of TMS were delivered to six different targets within the left dlPFC (Fig 1B). Targeting details have been reported previously (4). Briefly, the number of targets and spatial locations were chosen to balance the following: sampling of the anterior / posterior and medial / lateral planes in the middle frontal gyrus and sampling of the many utilized clinical dlPFC targets for depression (12). *t1* is close to the commonly used 5 cm average clinical target (10). *t2* corresponds to the EEG F3 site (11). *t3* is located midway between two targets: the BA9 definition (30) and a treatment site with resting fMRI activity negatively correlated to anterior subgenual cingulate cortex (sgACC) (10). *t4* corresponds to the treatment site with activity that is most negatively correlated to the sgACC (10). *t5* roughly corresponds to the TMS treatment which corresponded to better treatment success in the study by Herbsman *et al*. 2009 (13). *t6* roughly corresponds to the 5.5 cm average target (31). The six targets were registered to each individual brain using a non-linear inverse transformation from MNI space (32).

**Figure 1.**
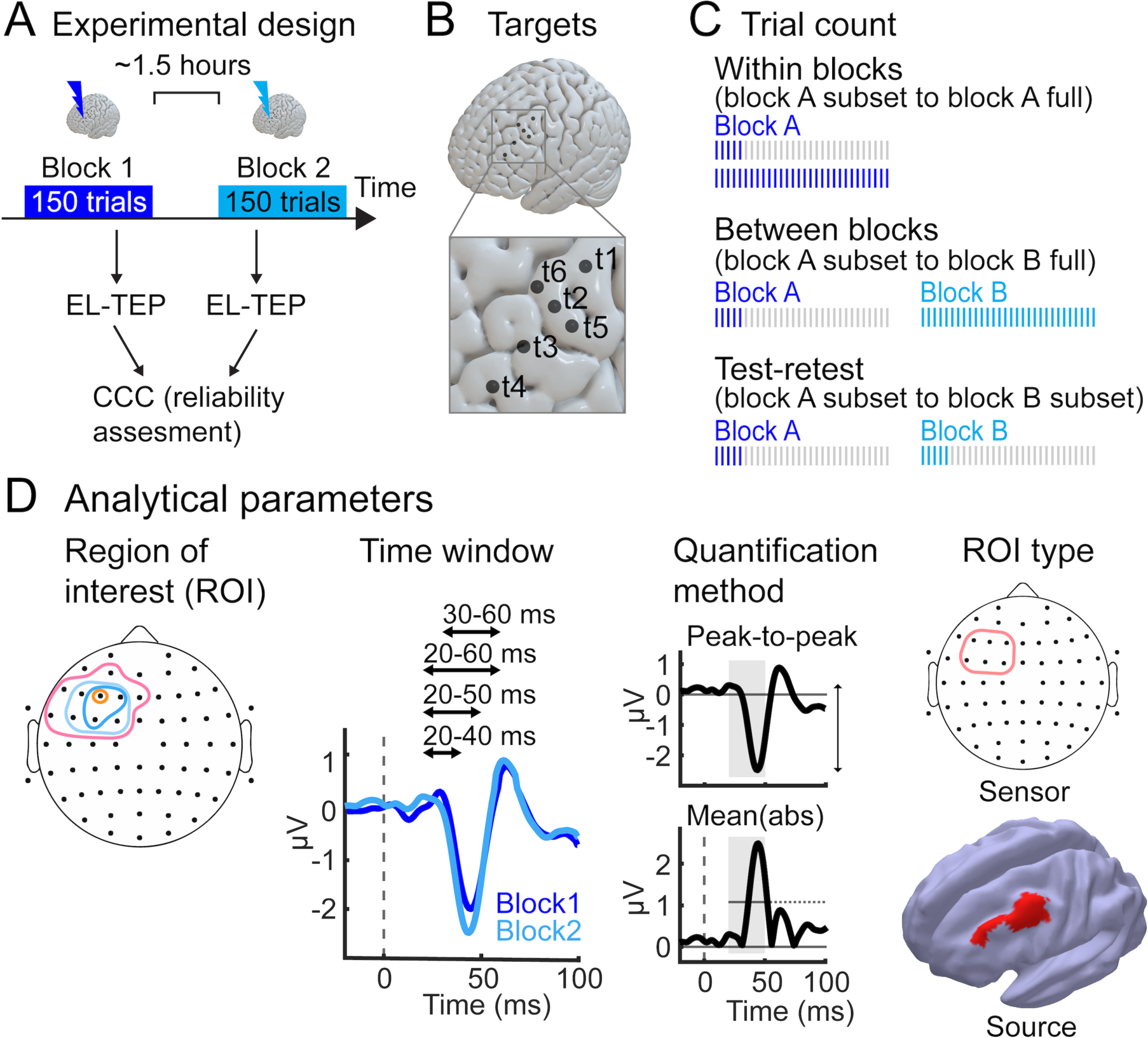
Study schematic. A) Experiment consisted of two single-pulse TMS-EEG blocks, separated by approximately 1.5 hours. For each subject, EL-TEPs were quantified for both experimental blocks. B) TMS was delivered to six different targets within the dlPFC. C) Retrospectively, each stimulation block was divided into subsets consisting of a smaller number of trials than the full sample (150 trials). Each subset was preprocessed separately. D) In addition to experimental manipulations (stimulation target and number of trials), we investigated how reliability of the dlPFC EL-TEP varied as a function of analytic parameters. Within the sensor space, four different ROI sizes and time windows, as well as two different quantification methods, were evaluated. Reliability of sensor space EL-TEPs was also compared to source space EL-TEPs.

#### 2.2.2. Experimental Design

The order of stimulation of the targets was pseudorandomized within the session. 150 single pulse TMS trials were collected for each experiment block. Two blocks of single pulse data were collected on the same day, separated by an average of 97 ∓ 37 minutes. TMS pulses were delivered every 1.5 s, jittered +/-25% (33). Work with intracranial stimulation has shown that an interstimulus interval of 1 s may not produce acute effects in cortico-cortical evoked potentials (34). To maintain levels of attention, subjects were shown neutral images on a screen and were instructed to keep eyes open during stimulation. Subjects were monitored throughout the experiment.

### 2.4. Electroencephalography

EEG was recorded using a 64-channel TMS-compatible amplifier (ActiCHamp Plus, Brain Products GmbH, Germany) with a 25 kHz sampling rate. Slim, freely rotatable, active electrodes (actiCAP slim, Brain Products GmbH, Germany) were used in a standard montage labeled according to the extended 10-20 international system. EEG data were online referenced to the ‘Cz’ electrode and recorded using BrainVision Recorder software (Brain Products GmbH, Germany). Impedances were monitored and percentage of channels with impedances <10 kΩ was 97.54 ∓ 2.31%. TMS “click” frequency matched noise masking and earmuffs, as described in detail previously (35), were applied to reduce off-target sensory effects.

#### 2.4.1. Preprocessing of TMS-EEG data

TMS-EEG preprocessing was performed with the fully automated AARATEP pipeline (version 2, (36)). Epochs were extracted from 800 ms before to 1000 ms after each TMS pulse. Data between 2 ms before to 12 ms after each pulse were replaced with values interpolated by autoregressive extrapolation and blending (36), downsampled to 1 kHz, and baseline-corrected based on mean values between 500 to 10 ms before the pulse. Epochs were then high-pass filtered above 1 Hz with a modified filtering approach to reduce the spread of pulse-related artifact into baseline time periods. Bad channels were rejected via quantified noise thresholds and replaced with spatially interpolated values (see (37) for all details on channel deletion and interpolation), with 1.8 +/-2.0% (mean +/-standard deviation) channels rejected on average. Eye blink artifacts were attenuated by a dedicated round of independent component analysis (ICA) and eye-specific component labeling and rejection using ICLabel (38), a modification from the original AARATEP pipeline introduced in version 2. An average of 1.7 +/-0.7 components were rejected at this stage. Various non-neuronal noise sources were attenuated with SOUND (39). Decay artifacts were reduced via a specialized decay fitting and removal procedure (36). Line noise was attenuated with a bandstop filter between 58-62 Hz. Additional artifacts were attenuated with a second stage of ICA and ICLabel labeling and rejection, with rejection criteria targeted at removing any clearly non-neural signals (see (37) for all data deletion criteria). 60.6% +/-10.3% of components were rejected, accounting for 30.3% +/-20.5% of variance in the signal at this stage. Data were again interpolated between -2 and 12 ms with autoregressive extrapolation and blending, low-pass filtered below 100 Hz, and average re-referenced. For complete details of the pipeline implementation, see (36) and source code at github.com/chriscline/AARATEPPipeline. For each target and analytical parameter combination (see section 2.5.), trial-averaged EL-TEPs were calculated with a 10% trimmed mean to prevent a small number of outlier trials from confounding results.

A subset of recorded conditions were processed two additional times with identical pipeline parameters as above to quantify variability between preprocessing runs due to non-deterministic parts of the pipeline, including random variation in ICA decomposition (40).

#### 2.4.2. Quantification of reliability

The ability of a biomarker to distinguish different individuals is best assessed by an intraclass correlation coefficient (ICC), which represents the ratio of the inter-participant variance to the total variance (41,42). To quantify within-session reliability of prefrontal EL-TEPs, we applied the concordance correlation coefficient (CCC), a form of ICC optimally modified to assess test-retest reliability (17,43), and defined the following correlation strength categories: low [<0.5], moderate [0.5, 0.7], and high [0.7, 1.0] correlation. For each target and analytical parameter combination, the magnitude of subjects’ EL-TEP of the first block was compared to that of the second block (Fig 1A) to obtain a group-level CCC value. Mean and 95% confidence intervals of the CCC were calculated across target and analytical parameter combinations. To compare CCCs across conditions, we quantified the means and ranges (Fig 2-3). Reliability calculations of different trial counts is described in section 2.4.3.

**Figure 2.**
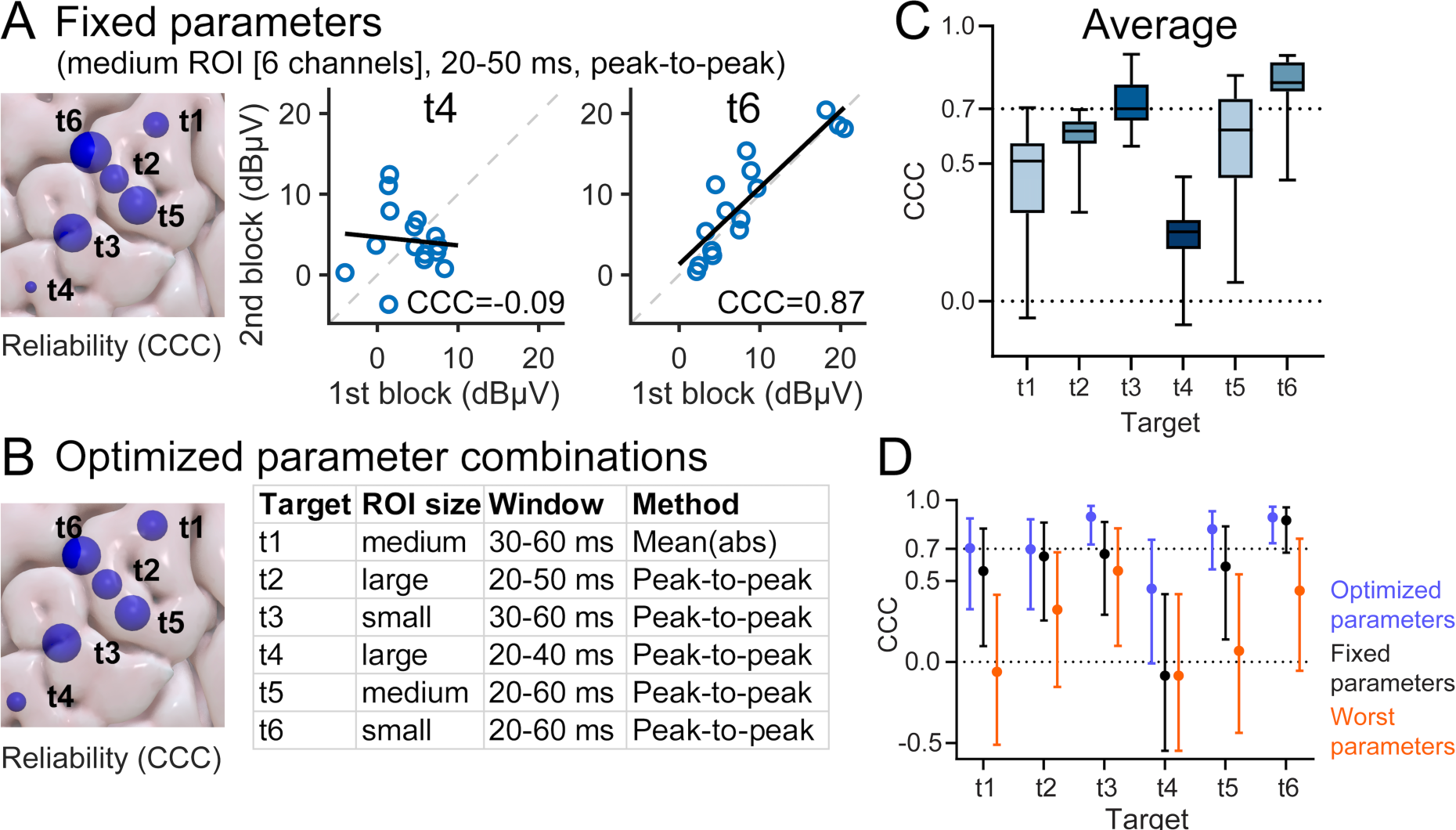
EL-TEP reliability across dlPFC targets for fixed and optimized analytic parameters. A) EL-TEP reliability across dlPFC targets using a fixed parameter combination previously used in an earlier study (medium ROI size, 20-50 ms time window, peak-to-peak amplitude; (4)). Scatter plots show EL-TEP amplitudes of the first and second experimental block for targets t4 and t6. B) Analytical parameter combinations that produced the most reliable EL-TEP amplitude for each target. On the left, marker size represents the CCC of the most reliable parameter combination. The table shows the most reliable parameters for each target. C) Average reliability across analytical parameters. D) Best and worst parameter combination compared to the fixed parameter combination for each target. In A and B, the blobs visualized on the brain surface represent CCCs but are not drawn to scale. In C, boxes represent the distribution of CCCs across analytical parameters in sensor space. In D, dots denote mean CCC and whiskers denote 95% CIs.

**Figure 3.**
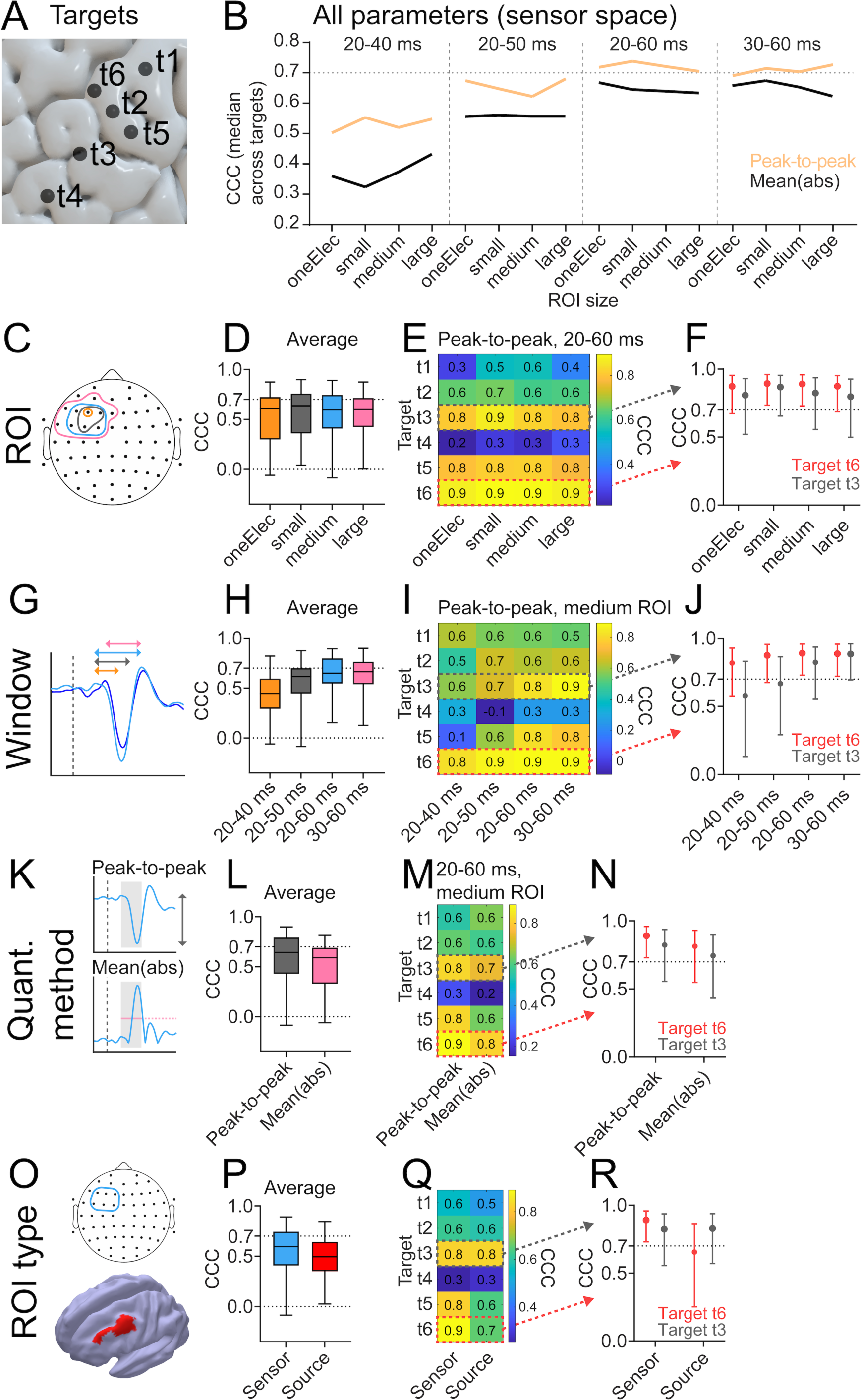
EL-TEP reliability across all analytical parameters. A) Stimulation targets in the dlPFC. B) Median CCCs across targets for all analytical parameters in sensor space. C-F) Effect of ROI size on EL-TEP reliability (in E and F peak-to-peak amplitude within 20-60 ms time window). G-J) Effect of time window on EL-TEP reliability (in I and J peak-to-peak amplitude within medium-size ROI). K-N) Effect of amplitude quantification method on EL-TEP reliability (in M and N medium-size ROI, 20-60 ms time window). B-N are in sensor space. O-R) Effect of ROI type (sensor versus source space) on EL-TEP reliability. In P-R the sensor space data represents the medium size ROI with peak-to-peak quantification and 20-60 ms time window. The box plots in D, H and L represent the distribution of CCCs across targets and other analytical parameters in sensor space. Last column (F, J, N and R) represents the two most reliable targets (t3 and t6) across the parameters (dots: mean, error bars: 95% CI). The dots denote mean CCC and whiskers denote 95% CIs.

In a supplementary analysis, we studied how reproducible each TEP is across the two experimental blocks. We calculated cosine similarity (44) across blocks for each stimulation target (Fig S1). Here, we did not restrict the analysis to any predetermined ROI but instead used all EEG channels as input. Cosine similarity was calculated within and between subjects.

#### 2.4.3. Subset analysis of TMS trial counts

In order to explore what the minimum number of trials were needed to obtain a reliable EL-TEP (CCC>0.7), 150-trial stimulation blocks were divided into 25, 50, 75, 100 and 125 trial subsets, including trials 1-25, 1-50, 1-75, 1-100, and 1-125. Each subset was preprocessed separately to simulate an experiment with a reduced number of trials, prior to EL-TEP amplitude quantification. EL-TEP was quantified using a fixed set of parameters based on our other CCC results which showed that a medium size ROI with six prefrontal channels, 20-60 ms time window, and peak-to-peak amplitude quantification is one of the most reliable parameter combinations (Fig 3). To obtain CCC, EL-TEP amplitude was compared either to 1) the amplitude of the same block when using a full set of 150 trials (*reliability within blocks*), 2) to the amplitude of another experimental block of the same target using a full set of 150 trials (*reliability between blocks*), or 3) to the amplitude of another block using the same amount of trials (*test-retest reliability*). *Reliability within blocks* was calculated to estimate the minimum number of trials needed to obtain a consistent EL-TEP response compared to the EL-TEP from a large number of trials, with minimal variation in coil positioning, brain state, or other experiment factors that may change over longer time scales. *Reliability between blocks* was calculated to evaluate the consistency of EL-TEPs over a longer time scale, varying just the number of trials in one block but using the maximum amount of available data (150 trials) in the other block as a proxy for a “ground truth” best TEP estimate. *Test-retest reliability* was performed to mimic experimental conditions in which one has less than 150 trials in each of both blocks, and provide an estimate of a minimum number of trials to obtain reliable EL-TEPs within a session.

#### 2.4.4 Quantification of within-block noise and SNR

As part of a supplemental analysis, we examined how within-block variance related to across-block reliability. We employed the bootstrapped standardized measurement error (bSME) (45) to quantify uncertainty in EL-TEP peak-to-peak and mean(abs) amplitude measures with 10000 bootstrap repeats per estimate, accounting for trial-to-trial variation in responses and number of trials aggregated. With bSME calculated on logarithmically-scaled dBμV values, we obtained bSME measures in units of dBμV, and calculated per-block SNR as the mean response amplitude (in dBμV) minus the estimated bSME value for the same block. Supplemental Fig S15 shows the results of analyses examining possible relationships between within-block variance and across-block reliability. While correlations between within-block response amplitude and within-block variance were identified (Fig S15 A, B, E, F), there were no strong correlations identified between within-block SNR and reliability across blocks (Fig S15 C, D, G, H).

### 2.5. Analytical parameters

Our goal was to characterize the reliability of prefrontal EL-TEPs across dlPFC stimulation targets and analytical parameters, defined as follows: quantification method, time window, ROIs size, and ROI type (Fig 1).

#### 2.5.1. Quantification method

We quantified EL-TEP amplitude by computing both a peak-to-peak amplitude metric and mean of the absolute amplitudes (mean(abs)), averaged across samples within each time window and ROI (see 2.5.2.). Because the number and timing of EL-TEP peaks from stimulation of the dlPFC are variable across targets and subjects (4,16), we quantified the peak-to-peak amplitude across different time windows using subtraction of the signal minimum from the signal maximum within a time window of interest (46), rather than the difference in amplitude of specific predefined TEP peaks. Both peak-to-peak and mean(abs) metrics were quantified in units of dBμV as linear-scale (μV) responses exhibited a roughly log-normal distribution (Fig S2).

#### 2.5.2. Time window

We focused on the early and local local response of the TEP (EL-TEP), which captures previously reported peaks (2,47,48) including the P30, N40, and P50, because 1) there is evidence for the neural basis of the early TEP components (34,49); 2) this time window is commonly studied in the literature (2,47,48); and 3) off-target sensory confounds are less prominent in these earlier components of the TEP (20,35,50–53). However, the optimal time window for quantifying prefrontal EL-TEPs remains unknown. Therefore, we assessed the test-retest reliability of EL-TEPs by calculating it within four distinct time windows: 20-40 ms, 20-50 ms, 30-60 ms, and 20-60 ms.

#### 2.5.3. ROI size (sensor space)

As there is no agreed upon optimal ROI size, we evaluated the effect of four predetermined prefrontal sensor-space ROIs on reliability (Fig 1D). These ROIs included a single electrode (F3), a small ROI (F3, F1, FC3), a medium ROI (F5, F3, F1, FC5, FC3, FC1), and a large ROI (F5, F3, F1, FC5, FC3, FC1, FT7, F7, AF3, Fz). We selected these ROIs to sample from and be centered over our dlPFC stimulation targets. As a followup analysis, to ensure these ROIs were not biasing results, we repeated the analyses with shifted ROIs (Fig S3 and Fig S4). See Supplementary Material for more details.

#### 2.5.4. Source space estimation

Subject-specific differences in gyral anatomy can cause underlying common cortical sources to project to the scalp in different topographies across subjects (54). To account for this and other related consequences of EEG volume conduction, we performed EEG source estimation and asked whether the source-space EL-TEPs would be more reliable compared to the sensor space TEPs. Using digitized electrode locations and individual head models constructed from subjects’ anatomical MRI data (55), subject-specific forward models of signal propagation from dipoles distributed over and oriented perpendicular to the cortical surface to electrodes on the scalp were constructed (56,57). Inverse kernels mapping measured scalp EEG activity to underlying cortical sources were estimated using weighted minimum-norm estimation (wMNE) as implemented in Brainstorm (58). A cortical region of interest (ROI) was constructed from a combination of HCP MMP1 atlas (59) parcels (46 L, 8Av L, and 8C L; Fig 1D lower right), and mapped from a common group template surface (ICBM152) to subject-specific cortical surfaces via surface-based morphometry.

## 3. Results

### 3.1. Reliability across stimulation targets

We first investigated the effect of EL-TEP reliability across different targets within the dlPFC (Fig 1B). We characterized the effect of region of interest (ROI) size and type, time window, and quantification method – together referred to as *analytic parameters* – on reliability. Across all analytic parameters in sensor space (ROI size, time window, and quantification method), targets t3 (mean CCC_t3_=0.72, range 0.56 to 0.90) and t6 (mean CCC_t6_=0.78, range 0.44 to 0.89) were highly reliable (CCC>0.7) on average, whereas all other targets had average CCCs under 0.7 (Fig 2C; CCC_t1_=0.44, CCC_t2_=0.59, CCC_t4_=0.24, CCC_t5_=0.56).

When using an *a priori* chosen fixed combination of parameters used previously to quantify the EL-TEP (medium sized sensor space ROI, 20-50 ms, peak-to-peak) (4), only target t6 was highly reliable (Fig 2A and D; CCC_t6_=0.87, CI [0.68, 0.96]). However, when considering the set of all tested analytic parameters in sensor space, all targets except t4 (CCC_t4_=0.62, CI [0.18, 0.85]) had a mean CCC of greater than 0.7 for at least one parameter combination (Fig 2B and D; CCC_t1_=0.70, CI [0.33, 0.89]; CCC_t2_=0.74, CI [0.42, 0.90]; CCC_t3_=0.98, CI [0.96, 0.99]; CCC_t5_=0.89, CI [0.70, 0.96]; CCC_t6_=0.95, CI [0.86, 0.98]). Of those, targets t3 and t6 had a lower 95% CI bound of greater than 0.7 indicating that at these two targets, with optimized parameters, almost all data should be expected to be highly reliable. Conversely, using a poor combination of parameters for a target could be expected to result in unreliable EL-TEPs; using the worst possible combination of analytic parameters (sensor space combination that yields the smallest CCC for each target) resulted in none of the targets being highly reliable (Fig 2D; all targets’ CCC < 0.6).

We also explored the spatial consistency of maximal EL-TEP amplitude across targets. When using a parameter combination of 20-60 ms time window, medium ROI and peak-to-peak quantification, 9/15 subjects exhibited the largest EL-TEP amplitude in the same target for both the first and second stimulation blocks. Moreover, 14/15 subjects exhibited the largest EL-TEP amplitude in either the same or adjacent stimulation target.

We next quantified cosine similarity across the two experimental blocks when all EEG channels were used (Fig S1). For similarity within subjects, no significant difference was found between the stimulation targets.

### 3.2. Reliability across analytical parameters

We next investigated the impact of various analytical parameters on the test-retest reliability of prefrontal EL-TEPs. Specifically, we examined the influence of ROI size and type, time window, and amplitude quantification method on the reliability. Table 1 shows the sensor space conditions where the lower bound of the 95% confidence interval for CCC exceeded 0.7, indicating a high level of reliability for most of the data. Of note, these reliable conditions used either the t6 or t3 targets and peak-to-peak quantification method. Median CCCs (across targets) for all analytical parameters in sensor space are shown in Fig 3B. CCCs for all targets and sensor-space parameters are shown in Fig S5 and Fig S6.

**Table 1.** Experimental conditions that yield highly reliable dlPFC EL-TEPs. The table shows sensor-space conditions in which the 95% confidence interval lower bound of the mean CCC was >0.7, indicating high reliability for most of the data.

| Target | ROI size | Time window | Quantification method | Mean CCC | 95% CI (lower) | 95% CI (higher) |
| --- | --- | --- | --- | --- | --- | --- |
| t6 | small | 20-60 ms | Peak-to-peak | 0.89 | 0.73 | 0.96 |
| t6 | medium | 20-60 ms | Peak-to-peak | 0.89 | 0.73 | 0.96 |
| t6 | medium | 30-60 ms | Peak-to-peak | 0.89 | 0.72 | 0.96 |
| t3 | small | 30-60 ms | Peak-to-peak | 0.90 | 0.73 | 0.96 |

#### 3.2.1. ROI size

We next examined the effect of sensor-space ROI size on prefrontal EL-TEP reliability (Fig 3C-F). We hypothesized that larger ROIs would be more reliable due to less susceptibility to the influence of artifact outliers in individual channels. The results showed that when averaging across all stimulation targets and time windows, EL-TEP reliability does not vary widely across ROIs tested (Fig 3D; *one electrode* mean CCC: 0.53, range -0.06 to 0.87; *small ROI* mean CCC: 0.56, range 0.040 to 0.90; *medium ROI* mean CCC: 0.55, range -0.085 to 0.89; *large ROI* mean CCC: 0.56, range 0.0023 to 0.87). When utilizing the most reliable time window (20-60 ms; Fig 3H) and amplitude quantification method (peak-to-peak; Fig 3L), both target t3 and target t6 demonstrated high reliability across various analytical parameters. However, in terms of the lower bound of the 95% confidence interval, only target t6 exhibited values exceeding 0.7, in small and medium ROIs (Fig 3F; CCC_small_=0.89, CI [0.73, 0.96]; CCC_medium_=0.89, CI [0.73, 0.96]). We next examined the influence of shifting ROIs anteriorly or posteriorly (Fig S3 and Fig S4). While other targets had stable CCCs across the shifted ROIs, for the most anterior and posterior targets (t1 and t4) the shift in ROI positioning had a large effect on the CCCs (Fig S4).

#### 3.2.2. Time window

Next, we studied the effect of the time window on prefrontal EL-TEP reliability in sensor space (Fig 3G-J and Fig S7-10). Because of numerous peaks previously reported within the EL-TEP time window (51,60,61), we hypothesized that a longer window (20-60 ms) that captured the P30/N40/P50 peaks would have greater reliability than shorter time windows (30-60 ms, 20-40 ms, 20-50 ms). When averaging across stimulation targets and ROI sizes, the longer time window (20-60 ms) did in fact have greater reliability (Fig 3H; mean CCC_20-60ms_=0.62, range 0.15 to 0.89) compared to the shorter 20-40 ms and 20-50 ms time windows (Fig 3H; mean CCC_20-40ms_=0.43, range -0.06 to 0.82; mean CCC_20-50ms_=0.55, range -0.085 to 0.87). However, one shorter time window that included later latencies, 30-60 ms, also had high reliability (Fig 3H; mean CCC_30-60ms_=0.61, range 0.13 to 0.90), suggesting that high EL-TEP reliability might be driven more by the later latencies of the EL-TEP rather than the length of the time window analyzed. When employing a medium ROI and peak-to-peak amplitude quantification, target t6 exhibited the greatest reliability, with CCC values exceeding 0.8 across all included time windows (Fig 3J; CCC_t6,20-40ms_=0.82, CI [0.58, 0.93]; CCC_t6,20-50ms_=0.87, CI [0.68, 0.96]; CCC_t6,20-60ms_=0.89, CI [0.73, 0.96]; CCC_t6,30-60ms_=0.89, CI [0.72, 0.96];). Target t3 demonstrated CCCs greater than 0.8 in the longer 20-60 ms and later 30-60 ms time windows (Fig 3J; CCC_t3,20-40ms_=0.58, CI [0.13, 0.83]; CCC_t3,20-50ms_=0.67, CI [0.29, 0.86]; CCC_t3,20-60ms_=0.82, CI [0.56, 0.94]; CCC_t3,30-60ms_=0.88, CI [0.69, 0.96];)). Peak-to-peak amplitudes for all time windows in medium size ROI are shown in Supplementary Material (Fig S7-10 for dBμV and Fig S11-14 for μV units). In summary, longer (20-60 ms) and later (30-60 ms) time windows increased EL-TEP reliability.

#### 3.2.3. Amplitude quantification method

We also studied the effect of EL-TEP quantification method on prefrontal EL-TEP reliability (in sensor space, Fig 3K-N). On average, peak-to-peak quantification was more reliable than mean(abs) (Fig 3L; CCC_peak-to-peak_=0.59, range -0.085 to 0.90; CCC_mean(abs)_: 0.52, range -0.060 to 0.81). When using a medium size ROI and 20-60 ms time window, both targets t6 and t3 had mean CCC>0.7 for both quantification methods (Fig 3N; CCC_t6,peak-to-peak_=0.89, CI [0.73, 0.96]; CCC_t6,mean(abs)_=0.81, CI [0.55, 0.93]; CCC_t3,peak-to-peak_=0.82, CI [0.56, 0.94]; CCC_t3,mean(abs)_=0.74, CI [0.43, 0.90]). In summary, peak-to-peak quantification increased reliability compared to the mean(abs).

#### 3.2.4. ROI type

We next compared the reliability of prefrontal EL-TEPs in sensor-space to those in source-space (ROI type; Fig 3O-R). We quantified the peak-to-peak EL-TEP amplitudes in sensor and source space and compared the reliability of dlPFC source-based ROI (see Methods) to a medium-size sensor space ROI. We hypothesized that because inverse estimation may better focus on the local stimulation response, source space analysis would yield more reliable EL-TEPs. In contrast, on average, EL-TEPs were more reliable in sensor space compared to source space (Fig 3P; mean CCC_sensor_=0.55, range -0.085 to 0.89; mean CCC_source_=0.49, range 0.026 to 0.85). When utilizing peak-to-peak quantification and a time window of 20-60 ms, both targets t6 and t3 exhibited mean CCC values exceeding 0.8 in sensor space (Fig 3R; CCC_t6,sensor_=0.89, CI [0.73, 0.96]; CCC_t3,sensor_=0.82, CI [0.56, 0.94]). However, in source space t6 displayed lower reliability compared to t3 (Fig 3R; CCC_t6,source_=0.65, CI [0.25, 0.86]; CCC_t3,source_=0.83, CI [0.57, 0.94]). In summary, EL-TEPs in sensor space (medium size ROI) were more reliable than our source-space EL-TEPs metric.

### 3.3. Influence of trial counts on EL-TEP reliability

Next, we explored how many single pulses of TMS (*trials*) are required to yield a reliable EL-TEP (Fig 4). We first calculated the CCCs to assess the reliability *within* blocks by comparing each *n*-trial subset to the full set of trials from the same block. This allowed us to compare trial subsets to a full “gold standard set” of trials, answering the question *how close to the full EL-TEP block can one get from a subset of trials*? We found that for all targets except t4, 50 trials were enough to produce similar EL-TEPs to the full block (Fig 4C-D; CCCs > 0.7). For targets t3 and t6, only 25 trials were needed to yield high reliability compared to the full block (CCC_t3,within,25_=0.74; CCC_t6,within,25_=0.86).

**Figure 4.**
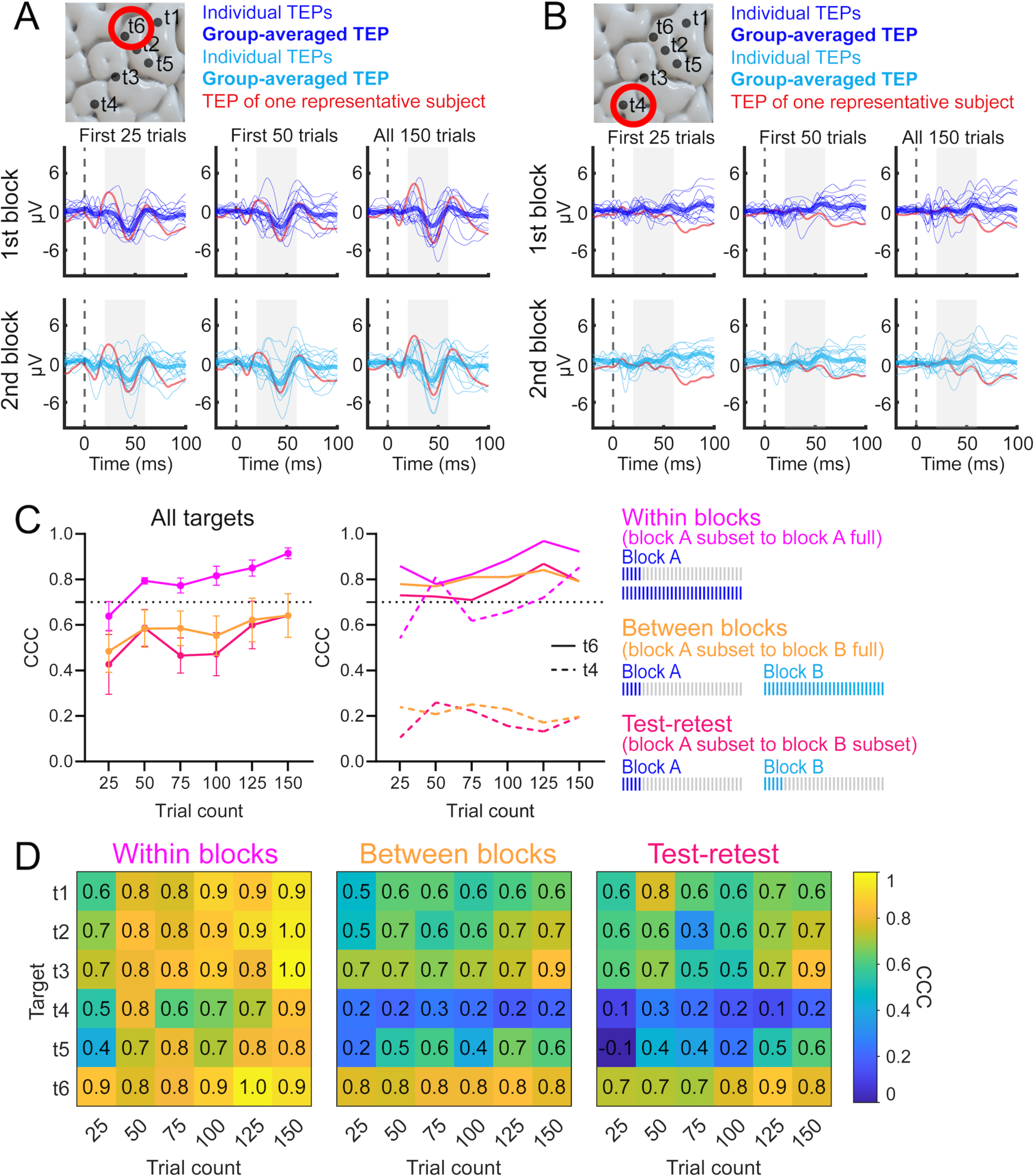
Effects of trial count on EL-TEP reliability. A-B) Each thin trace represents the TEP of one subject, averaged over electrodes of a medium size ROI. Thick traces represent the average TEP across subjects. Crimson color highlights TEPs from one representative subject. A) Example TEP traces for a reliable target (t6). B) Example TEPs for a non-reliable target (t4). TEPs for other targets are shown in Fig S17. C) Number of trials needed to produce reliable EL-TEPs. Left subplot shows the average CCC across dlPFC targets and the right subplot shows targets t6 and t4. Blue lines represent reliability within blocks, i.e. each subset of trials is compared to the full sample of 150 trials of the same block. Orange lines represent reliability between blocks, i.e. each subset of trials is compared to the full sample of 150 trials of a different block. Pink lines represent the test-retest reliability, i.e. each subset compared to the same subset of a different block. In the left subplot the dots denote mean CCC across targets ±SEM. D) Within-block, between-block and test-retest reliability across trial count and target.

We then calculated the reliability *between* blocks to evaluate the consistency of EL-TEPs over a longer time scale (∼1.5 hours). To do so, we studied the effect of the number of trials in the first block against the maximum amount of available data (150 trials) in the other block as a proxy for a “ground truth” best TEP estimate (Fig 4 C-D). For target t6, only 25 trials were enough to yield high reliability between blocks (CCC_t6,between,25_=0.78). For targets t1, t2 and t3, subsets of 50 or more trials produced moderate to high reliability (CCC_t1,between,50_=0.63; CCC_t2,between,50_=0.65; CCC_t3,between,50_=0.71). Target t4 yielded low reliability between blocks regardless of the trial count (CCC < 0.3).

Next, to mimic experimental conditions in which one may record fewer than 150 trials per block, we evaluated the *test-retest* reliability of experimental blocks by comparing the amplitude of one block to the amplitude of another block using the same number of trials (Fig 4C-D). For target t6, only 25 trials were enough to produce high test-retest reliability (CCC_t6,test-retest,25_=0.73). Target t3 produced high and t5 produced moderate test-retest reliability with more than 125 trials (CCC_t3,test-retest,125_=0.72; CCC_t5,test-retest,125_=0.52). Target t4 produced low test-retest reliability regardless of the trial count (CCC < 0.3).

In summary, when stimulating at targets t3 or t6, as few as 25 trials were enough to yield EL-TEPs that were reliable within an experimental block. Moreover, for target t6 as few as 25 trials were also sufficient for high test-retest and between-block reliability. For other dlPFC targets, reliable EL-TEPs between blocks may require more trials.

## 4. Discussion

### Summary of results

In the current study, we investigated how reliability of the prefrontal EL-TEP varied as a function of both stimulation target and analytic parameters. We applied single pulse TMS to six clinically-relevant targets within the dlPFC. We hypothesized that due to larger muscle artifacts observed after stimulation of more anterior targets (4), EL-TEPs would be more reliable after stimulation of posterior and medial dlPFC targets. Regarding the effect of analytic parameters (region of interest, time window, quantification method, trial count) on EL-TEP reliability, we hypothesized that later time windows within the early TEP, larger ROIs, source-space TEPs rather than sensor-space TEPs, and more trials would all increase reliability. Our main findings are summarized as follows: 1) The medial dlPFC TMS target was most reliable, while the most anterior dlPFC target was least reliable (Fig 2C); 2) targets in both the anterior and posterior dlPFC were reliable after optimizing analysis parameters (Fig 2D); 3) analyzing EL-TEPs using peak-to-peak quantification, longer and later time windows, and sensor-space EL-TEPs strengthened EL-TEP reliability (Fig 3); 4) ROI size did not appear to have a strong influence on EL-TEP reliability; 5) reliable EL-TEPs can be achieved with only 25 trials when stimulating the medial dlPFC. Together, these findings highlight that, a) EL-TEPs can be reliable, particularly if stimulating a medial prefrontal target; b) there is an urgent need to fine-tune analytic parameters to optimize reliability; c) ROI size used for analysis may not be as important for achieving a reliable EL-TEP biomarker; and d) the number of trials needed for obtaining a reliable EL-TEP may not be as high as previously assumed.

### Effect of stimulation target on reliability

How reliable are TMS-EEG measurements of excitability (EL-TEP) when stimulating the dlPFC? Our current findings underscore the significance of target selection, as it emerged as a crucial factor to obtaining highly reliable EL-TEPs. Specifically, we observed that with a medial target (t6), reliable EL-TEPs were obtained regardless of the chosen ROI size and time window (CCC from 0.72 to 0.89 for sensor-space peak-to-peak quantification). The most anterior site (t4) exhibited unreliable EL-TEPs irrespective of ROI size and time window (CCC from -0.085 to 0.45 for sensor-space peak-to-peak quantification), but another anterior (albeit less so) site (t3) exhibited reliable EL-TEPs for some parameter combinations (CCC from 0.56 to 0.90 for sensor-space peak-to-peak quantification). To our knowledge this is the first study evaluating differences in reliability of TEP across dlPFC subregions. There are a few earlier prefrontal TEP reliability studies, however. Kerwin *et al.* (17) reported low reliability of CCC < 0.6 for early TEPs (N40 peak) across blocks in a given day. Lioumis *et al.* (47) reported high Pearson correlations of between 0.88 to 0.96 for early dlPFC TEP peaks across separate days. These inconsistencies across the two papers may be due to several factors, including differences in noise masking, which is now common in the field (53), and differences in pre- and post-processing. In the current study, we used the newest sensory masking technique which involves auditory masking, foam separator between TMS coil and the scalp and over-the-ear protection (62). Future TEP studies should continue to apply the newest experimental and analytic techniques to reduce off-target sensory effects and continue to optimize EL-TEP reliability by exploring the effect of EEG amplifier, TMS coil type, pulse waveform, stimulation intensity, variations in TMS-EEG preprocessing, demographics (15). Finally, it is important to systematically investigate the reliability of EL-TEPs in other brain regions outside of dlPFC. In summary, we observed higher within-session EL-TEP reliability after TMS to the medial dlPFC, and that other dlPFC targets can produce reliable EL-TEPs after appropriate analytic considerations.

### Relationship between EL-TEP *amplitude, SNR,* and *reliability* across stimulation targets

We recently reported that posteromedial targets exhibited larger *amplitude* EL-TEPs compared to anteromedial sites (4). We would like to point out that *amplitude* does not necessarily co-fluctuate with *reliability*. Case in point, while our previous work demonstrated anterior dlPFC targets to elicit smaller amplitude EL-TEP responses (4), in this study we report some anterior sites with low reliability (t4) and others with high reliability (t3). Thus, both amplitude and reliability should be assessed when considering applying it to study the neural effect of an intervention (15,63–66).

We also hypothesized that within-block signal-to-noise would positively correlate with across-block reliability. However, our supplemental analysis did not reveal strong trends supporting this hypothesis (Fig S15), suggesting that factors contributing to within-block variance (e.g. trial-to-trial noise in EEG signals) may not be the main factors contributing to across-block degradation in reliability. Our trial subset analyses (Fig 4) indicating high within-block reliability in EL-TEP measures with just 25-50 trials further support this conclusion. Instead, longer time scale factors such as slow changes in brain state, drifts in stimulation coil positioning, or changes in electrode impedances may be more dominant factors and warrant further investigation in future experiments focused on improving EL-TEP reliability.

### Contribution of analytic parameters across stimulation targets

We observed that the size of the grouping of prefrontal electrodes used for analysis (ROI) did not significantly impact reliability of the EL-TEP (Fig 3C-F). We found that, on the contrary, the time window used in the analysis did impact reliability: earlier / narrower time windows (20-40 ms, 20-50 ms) produced EL-TEPs with lower reliability for all targets except t1 (most posterior target), whereas later (30-60 ms) and longer windows (20-60 ms) produced EL-TEPs with higher reliability (Fig 3G-J). The choice of amplitude quantification method also influenced reliability: peak-to-peak was more reliable than the mean of the absolute value (Fig 3K-N), although both had moderate reliability. Our metric for absolute amplitude, the mean of the absolute values within a time window, is equivalent to the common area-under-the-curve (AUC) metrics of TEP amplitude (7,35,67), except with a normalization for window length. Our method for peak-to-peak calculation, similar to that used in (68), differs from some other approaches that rely on temporal peak detection (e.g. (69,70)). Therefore, it should not be assumed that our reported reliability of peak-to-peak amplitude would be equal to different peak-to-peak metrics used in other EEG studies. When specific EL-TEP peaks are more well-defined, other methods for peak-to-peak calculation should be examined for reliability. Finally, we found that source localized EL-TEPs were less reliable than sensor-space EL-TEPs (Fig 3O-R). It is worth noting that for targets such as t3 (anterior target) where EL-TEP amplitude is low, the choice of quantification parameters may substantially impact reliability. Notably, the optimal parameter combination for achieving higher reliability varied for each target, as illustrated in Figure 2B. If researchers opt for a fixed parameter set across multiple stimulation targets, based on our results we recommend utilizing peak-to-peak amplitude quantification with a time window of either 20-60 ms or 30-60 ms in sensor space to maximize reliability (Fig 3B). However, from this work we find that to maximize target-specific reliability for EL-TEPs, each target may require different analytic parameters. When evaluating new stimulation targets, a distinct early data collection stage focused on reliability testing will help inform choice of optimal stimulation target to maximize EL-TEP reliability. This separate, early stage reliability evaluation could also be used to optimize and fix analytic parameters prior to primary data collection, allowing the opportunity to follow best practices in brain imaging research including pre-registering details of study design and outcome metrics.

### Influence of trial number on reliability

Here, we investigated the number of single pulses of TMS (trials) required to obtain a reliable prefrontal EL-TEP. To address this, we subdivided the data into different sets of *n*-trials and compared reliability across resulting EL-TEPs. An important note about our methods is that we preprocessed each set of *n*-trials separately in order to mimic the preprocessing that would be done in an experiment that only collected *n* number of trials. We found that for all except the most anterior dlPFC target (t4), averaging only 50 trials to compute the EL-TEP yielded a similar EL-TEP response compared to the EL-TEP from 150 trials (reliability *within* blocks; Fig 4D). Moreover, for three of the targets (t1, t2 and t3), a moderate to high consistency of EL-TEPs over a longer time scale (reliability *between* blocks) was obtained with only 50 trials (Fig 4D). When comparing less than 150 trials across both blocks (*test-retest* reliability), two targets produced high to moderate reliability with 125 trials. Notably, for the most medial target (t6), as few as 25 trials were sufficient to obtain high reliability both within and between blocks, as well as high test-retest reliability (Fig 4A, C and D). For the least reliable target (t4), reliability between blocks and test-retest reliability were low regardless of the trial count (Fig 4B, C and D).

These results are somewhat inconsistent with Kerwin *et al*. (17) which reported that the early TEP (in this case, the ‘N40’) was only moderately reliable (CCC ∼ 0.5) across experimental blocks, regardless of how many trials were used. However, several important differences between this study and (17) could account for this inconsistency: different EEG recording systems, different preprocessing pipelines, and slightly different stimulation targets. On the other hand, our results are compatible with the supplementary results described by Casarotto *et al*. (64). They estimated the minimum amplitude of the EL-TEPs that can be recovered from baseline EEG after averaging a certain number of trials, and the minimum number of trials required for signal reliability. They found that, if data has a good signal-to-noise ratio, as low as 20 trials are enough. Our results are also consistent with Lioumis *et al.* (47) in showing high reliability of the EL-TEP, though in that study they only examined 100 trials per condition and did not investigate what the minimum number of trials should be to obtain a reliable EL-TEP. Moreover, they used a different way of quantifying EL-TEPs compared to the current study. We recommend that future studies aiming to achieve reliable EL-TEPs from stimulation of the dlPFC should collect at least 50 trials per target. If requiring 25 or fewer trials, for example for real-time monitoring and closed-loop adaptive stimulation, our data suggest that the t6 target (in MNI coordinates x=-32, y=26, z=52) would be the optimal target. Regardless, reliability testing should be performed when using other equipment and experimental and analytic procedures.

### Limitations

While this study presents several important findings, numerous points limit generalizability and warrant careful consideration. First, the reliability assessment was limited to within-session measurements, as data between different days were not collected in this study. Within-session reliability is useful to determine how stable EL-TEPs are prior to an intervention, and as such, how confident one can be that a pre/post-intervention change *on a single day* is attributed to the intervention. For longitudinal studies, this type of test-retest reliability investigation must be performed across days. Second, it is important to note that a highly reliable *signal* does not necessarily imply high *biological validity* (15). Coupling these noninvasive studies with invasive investigations (5,34,49) to determine the neural response contributing to the EL-TEP will help reveal whether the EL-TEP is *biologically valid*. Third, details of the TMS-EEG preprocessing pipeline can affect reliability, and future improvements in preprocessing methods may increase reliability in a manner that reduces sensitivity to a specific stimulation target or analytic parameters (Fig S16). Fourth, our results may not generalize to other TEP quantification metrics, such as methods based on peak detection (16) or other time windows. Fifth, this study primarily focused on a predetermined set of analytic parameters. Future work should explore reliability across a broader range of parameter combinations, encompassing more refined (and potentially personalized) time windows, an expanded (and potentially personalized) set of sensor-space and source-space ROIs, and features in the spectral domain, which could further enhance reliability. Finally, it is important to acknowledge the influence of the experimental setup and hardware employed in this study, including the TMS coil, EEG system, and the use of a robotic arm for accurate targeting. Consequently, even if the dlPFC targets are the same as those reported here, for each hardware/software setup it may be beneficial for researchers to perform a reliability analysis to fine-tune experimental and analytic parameters and maximize TEP reliability (15).

### Conclusions

We characterized the test-retest reliability of the EL-TEP after stimulation of different dlPFC subregions. We found that EL-TEP reliability was sensitive to stimulation target in the dlPFC, with medial and anterior targets producing the most and least reliable EL-TEPs, respectively. The EL-TEP was sensitive to the quantification method, where peak-to-peak measurement in the 20-60 ms time window measured in sensor space typically provided the most reliable EL-TEPs. Finally, reliable EL-TEPs were produced after only 25 single TMS pulses to the medial dlPFC. These results have important implications for the EL-TEP as a measure of cortical excitability and potential clinical biomarker.

## Acknowledgements

We extend our deep gratitude to all of our research participants. We would also like to acknowledge the generous contributions of the members of the Personalized Neurotherapeutics Laboratory for helpful feedback on the manuscript and throughout the course of the study. We acknowledge Recep Ozdemir for his valuable assistance in implementing the original cosine similarity code adapted for this study. This research was supported by the National Institute of Mental Health under award number R01MH126639, R01MH129018, and a Burroughs Wellcome Fund Career Award for Medical Scientists (CJK). JG was supported by personal grants from Orion Research Foundation, the Finnish Medical Foundation and Emil Aaltonen Foundation. JMR was supported by the Department of Veterans Affairs Office of Academic Affiliations Advanced Fellowship Program in Mental Illness Research and Treatment, the Medical Research Service of the Veterans Affairs Palo Alto Health Care System and the Department of Veterans Affairs Sierra-Pacific Data Science Fellowship.

## Supplementary Material

**Figure S1.**
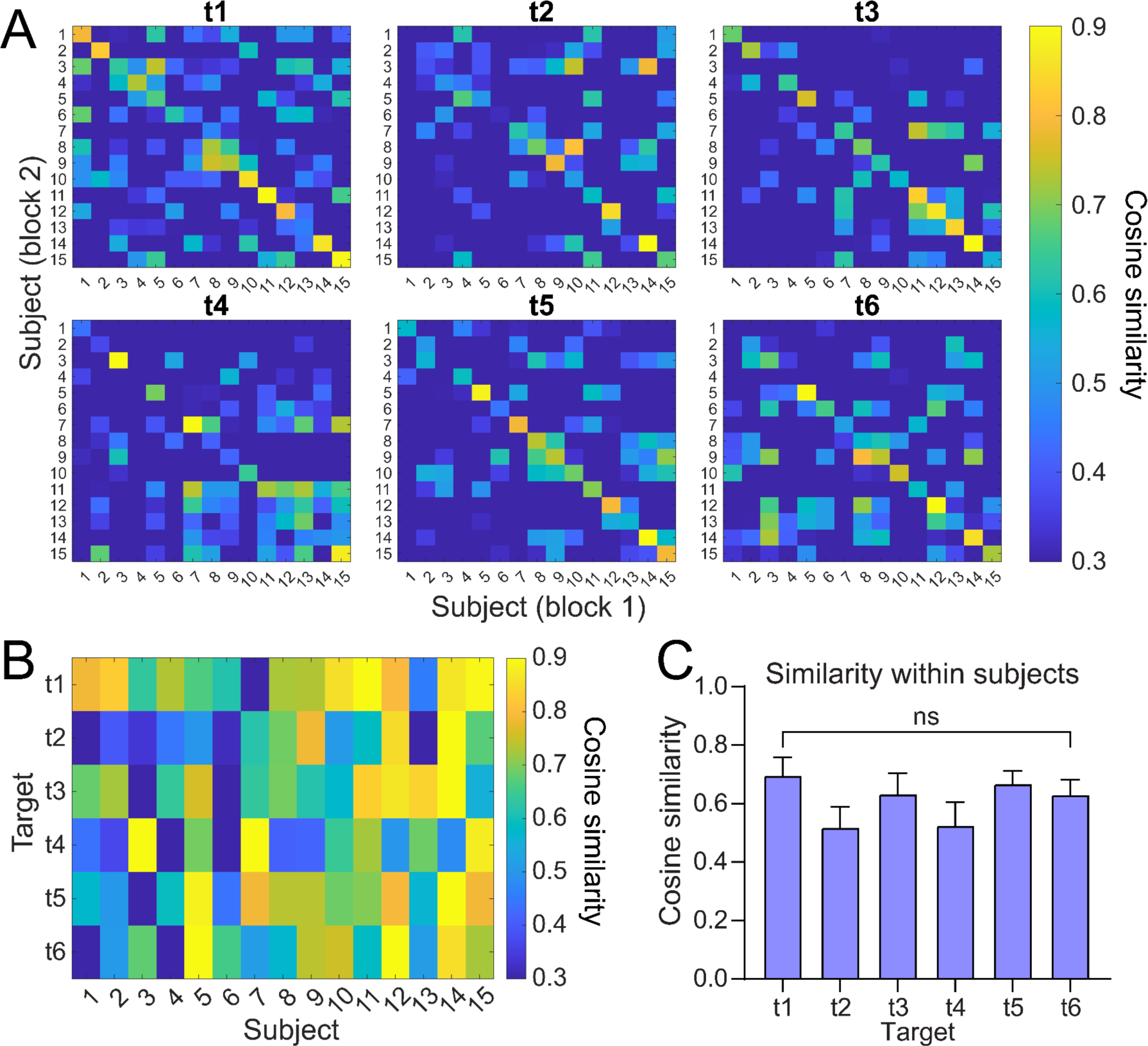
Cosine similarity of EL-TEPs. Cosine similarity comparing the first to the second block, calculated within a 20-60 ms time window using all EEG channels. A) Cosine similarity across all subjects. B) Cosine similarity within subjects. C) Within-subject similarity for each target. Repeated measures ANOVA showed no significant differences between the stimulation targets (*F*_3.33,46.65_=1.75, *P*=0.16). In C the bars denote mean cosine similarity across subjects ±SEM.

**Fig S2.**
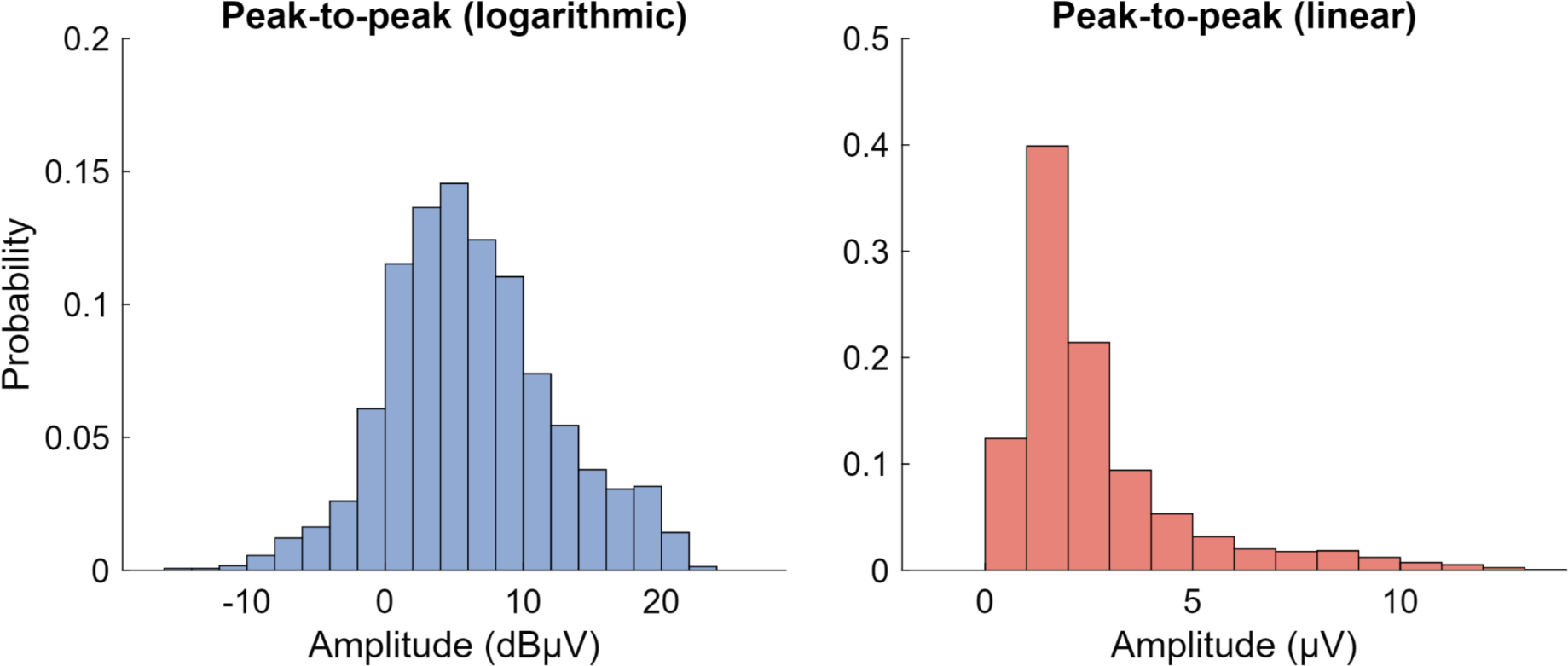
Histograms of sensor space peak-to-peak EL-TEP amplitudes in logarithmic (dBμV) and linear (μV) values. Logarithmic distribution passed D’Agnostino & Pearson normality test (*P*=0.23), whereas the linear distribution did not (*P*<0.0001). Moreover, the linear distribution passed D’Agnostino & Pearson lognormality test (*P*=0.23)

**Figure S3.**
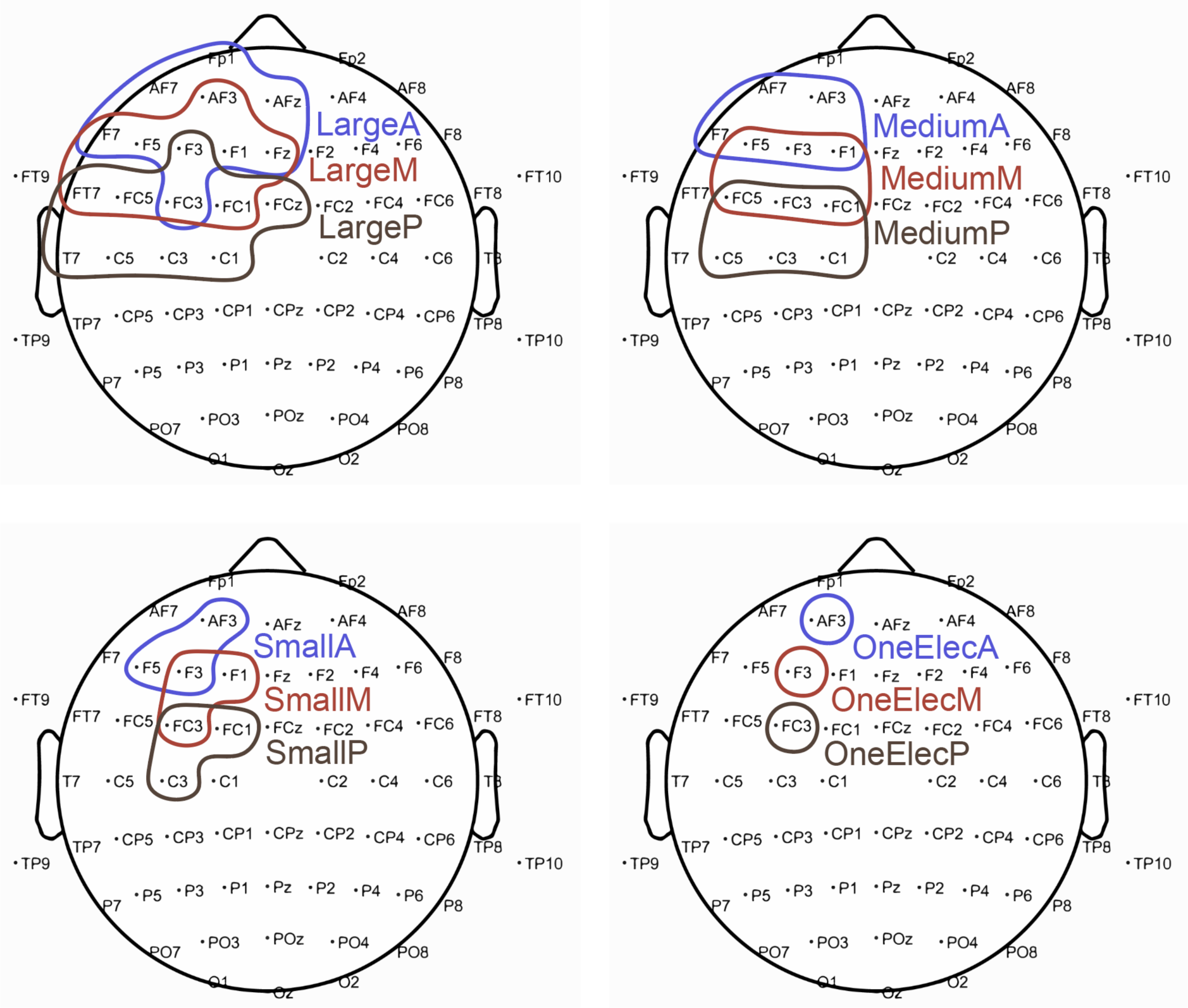
Locations of the shifted ROIs.

**Figure S4.**
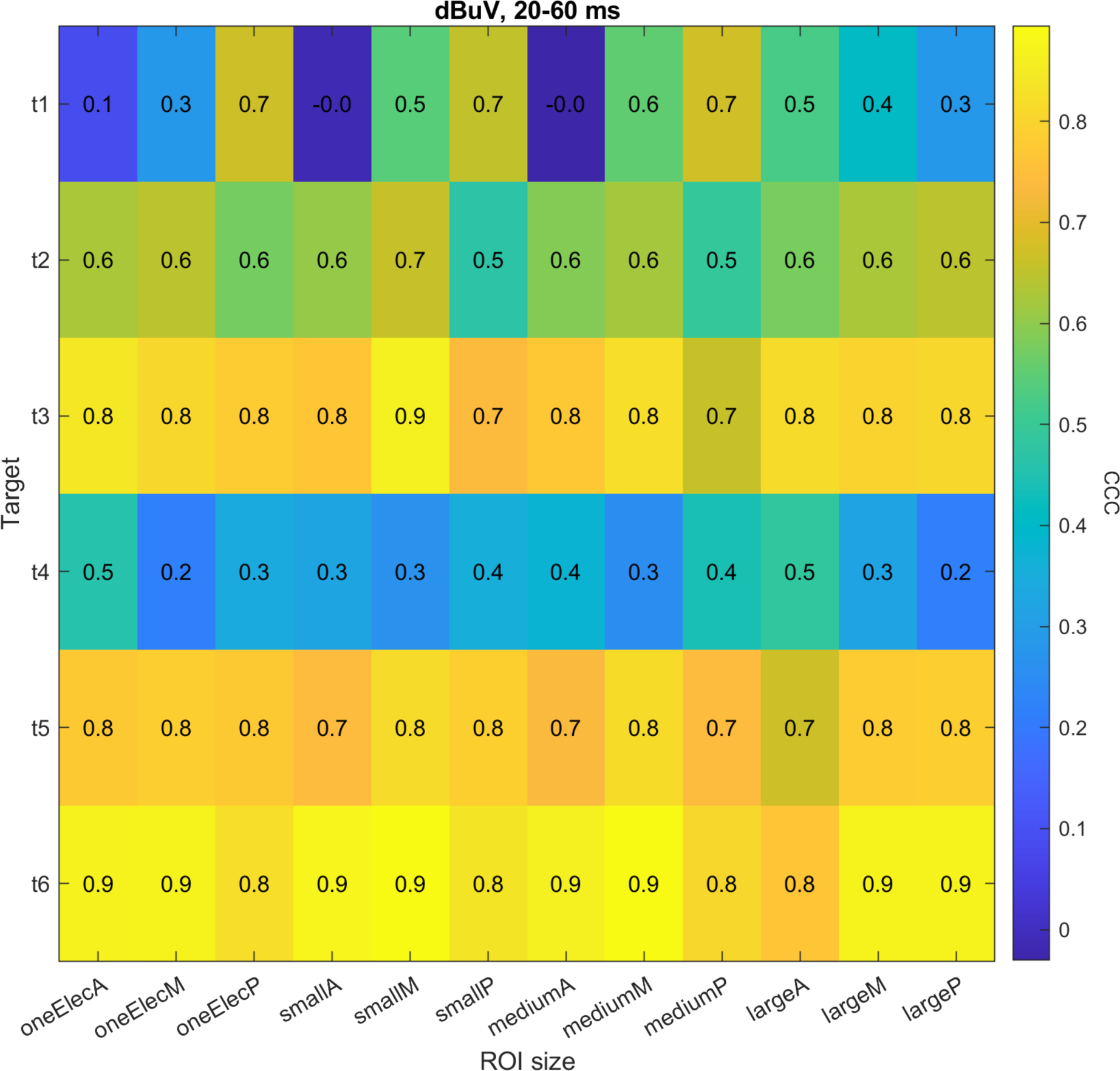
Effect of shifting the ROI on EL-TEP reliability. To examine whether the spatial positioning of the ROIs affects the results, we additionally shifted each ROIs in AP direction (three different positions for each ROI size) and extracted the TEPs and CCCs for these shifted ROIs. The heatmap shows CCCs for all shifted ROIs (peak-to-peak amplitude, 20-60 ms time window).

**Fig S5.**
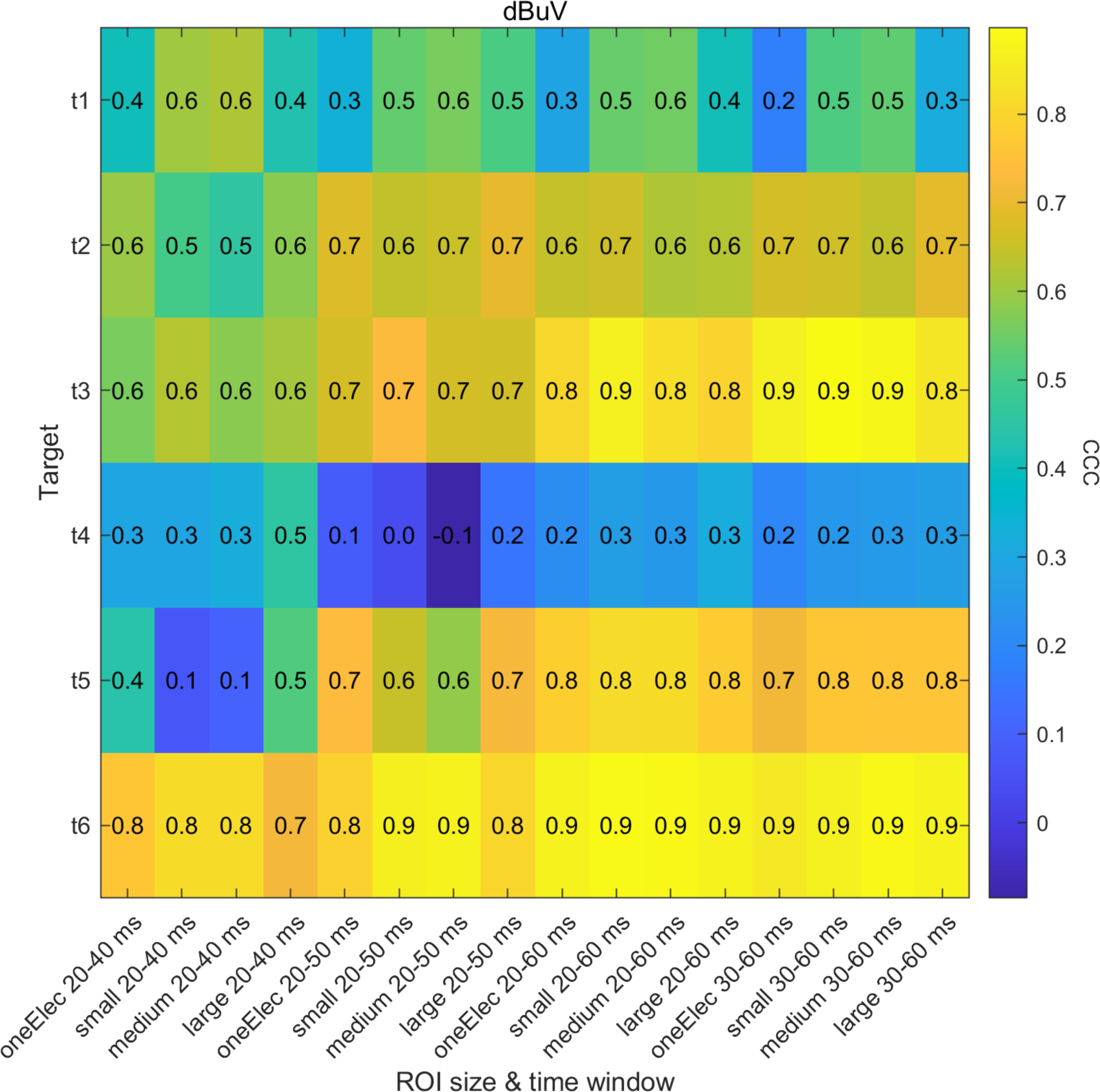
Effect of time window and ROI size on reliability for peak-to-peak quantification (in dB units).

**Fig S6.**
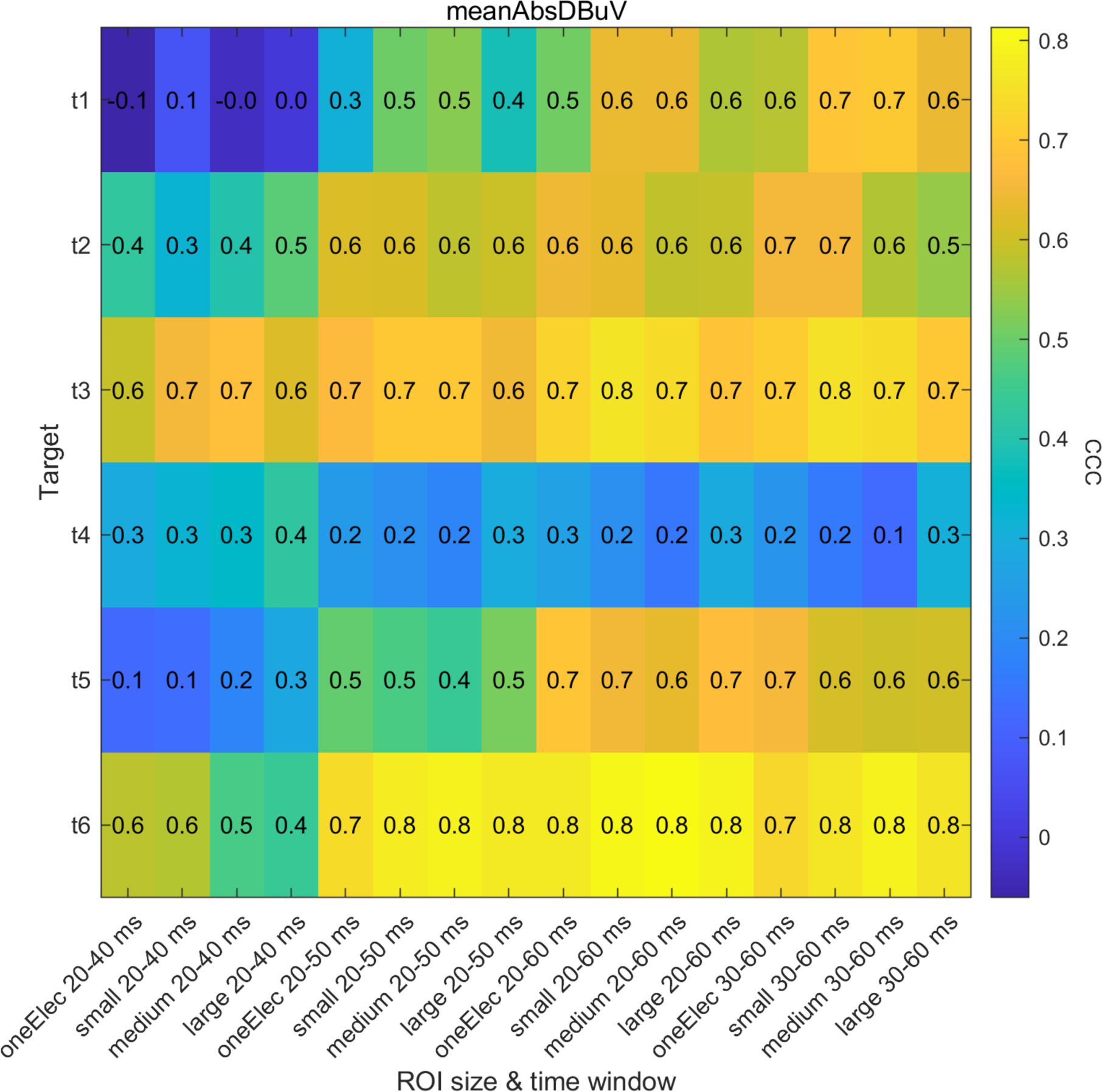
Effect of time window and ROI size on reliability for mean(abs) quantification (in dB units).

**Figure S7.**
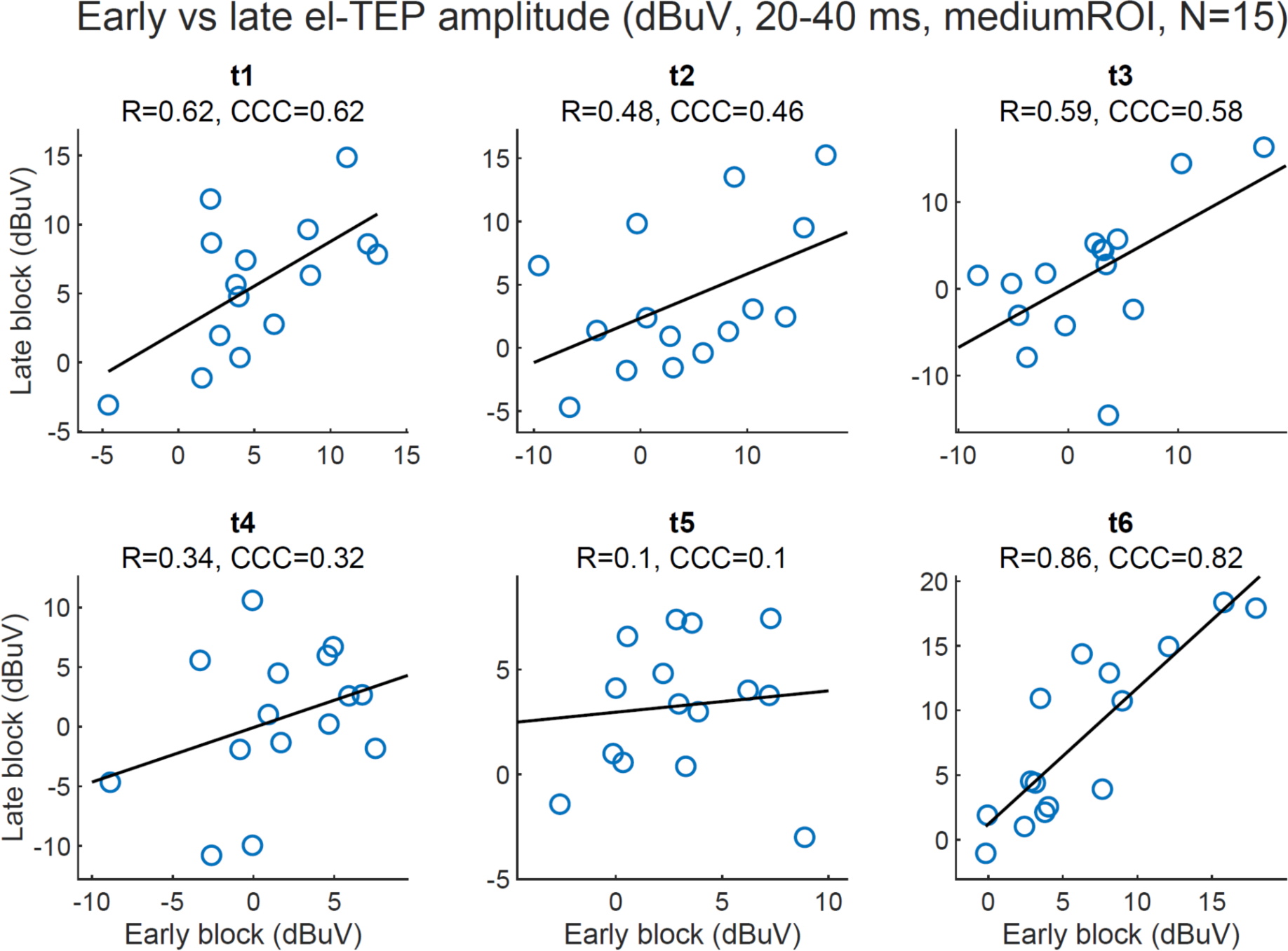
EL-TEP amplitudes (medium ROI, 20-40 ms time window, peak-to-peak) of the first and the second experimental block. Each dot represents one subject (N=15).

**Figure S8.**
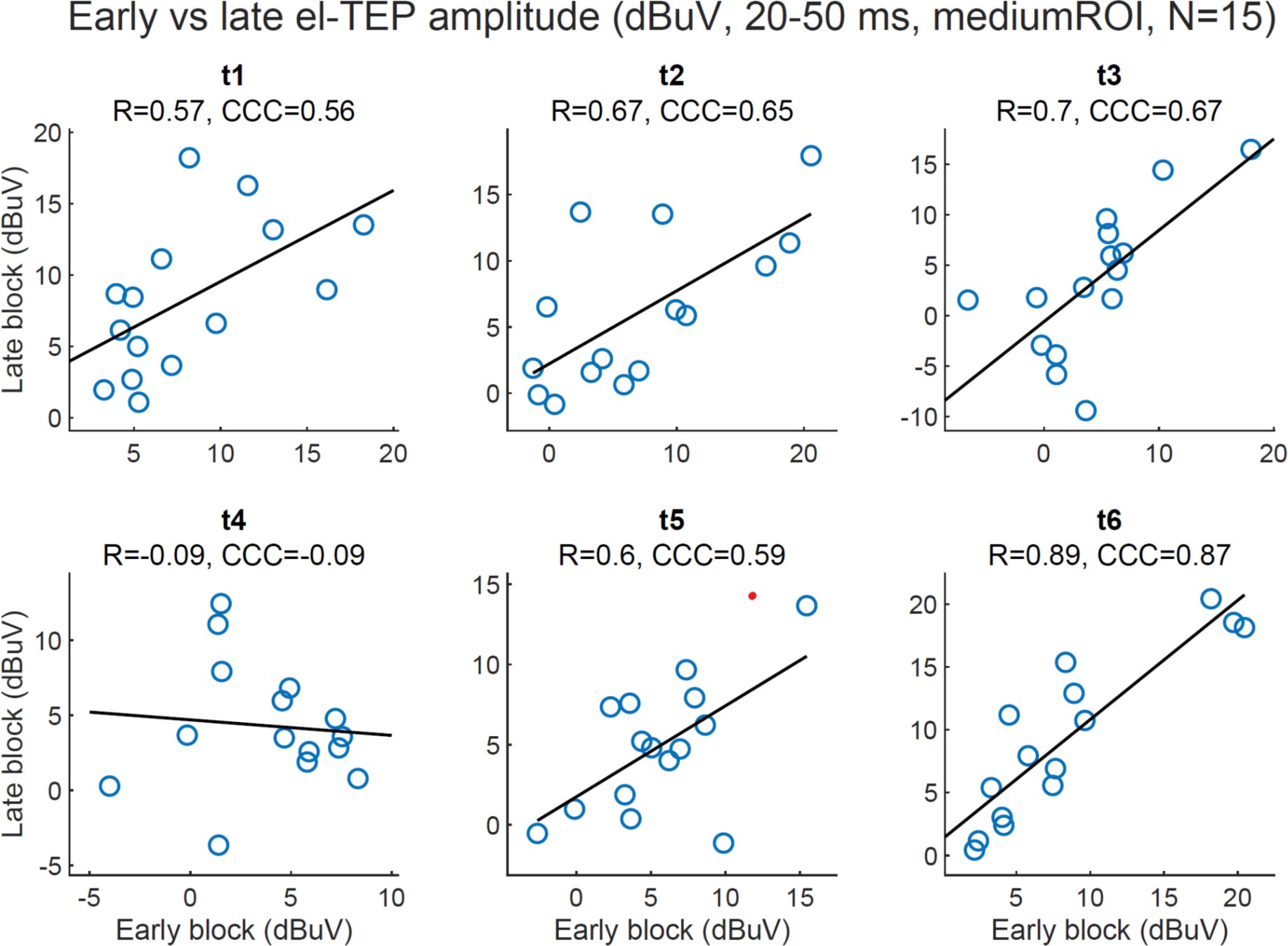
EL-TEP amplitudes (medium ROI, 20-50 ms time window, peak-to-peak) of the first and the second experimental block. Each dot represents one subject (N=15).

**Figure S9.**
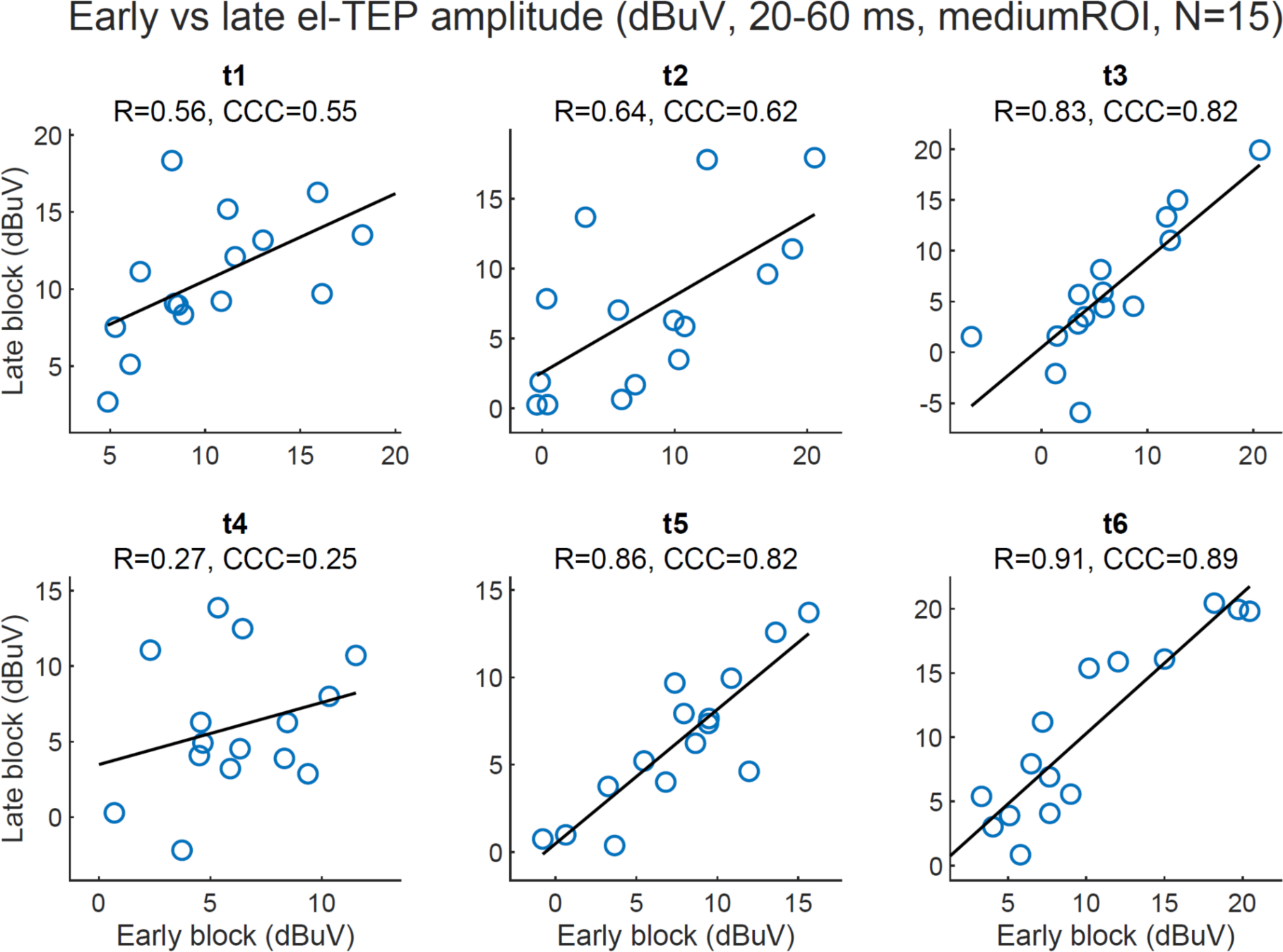
EL-TEP amplitudes (medium ROI, 20-60 ms time window, peak-to-peak) of the first and the second experimental block. Each dot represents one subject (N=15).

**Figure S10.**
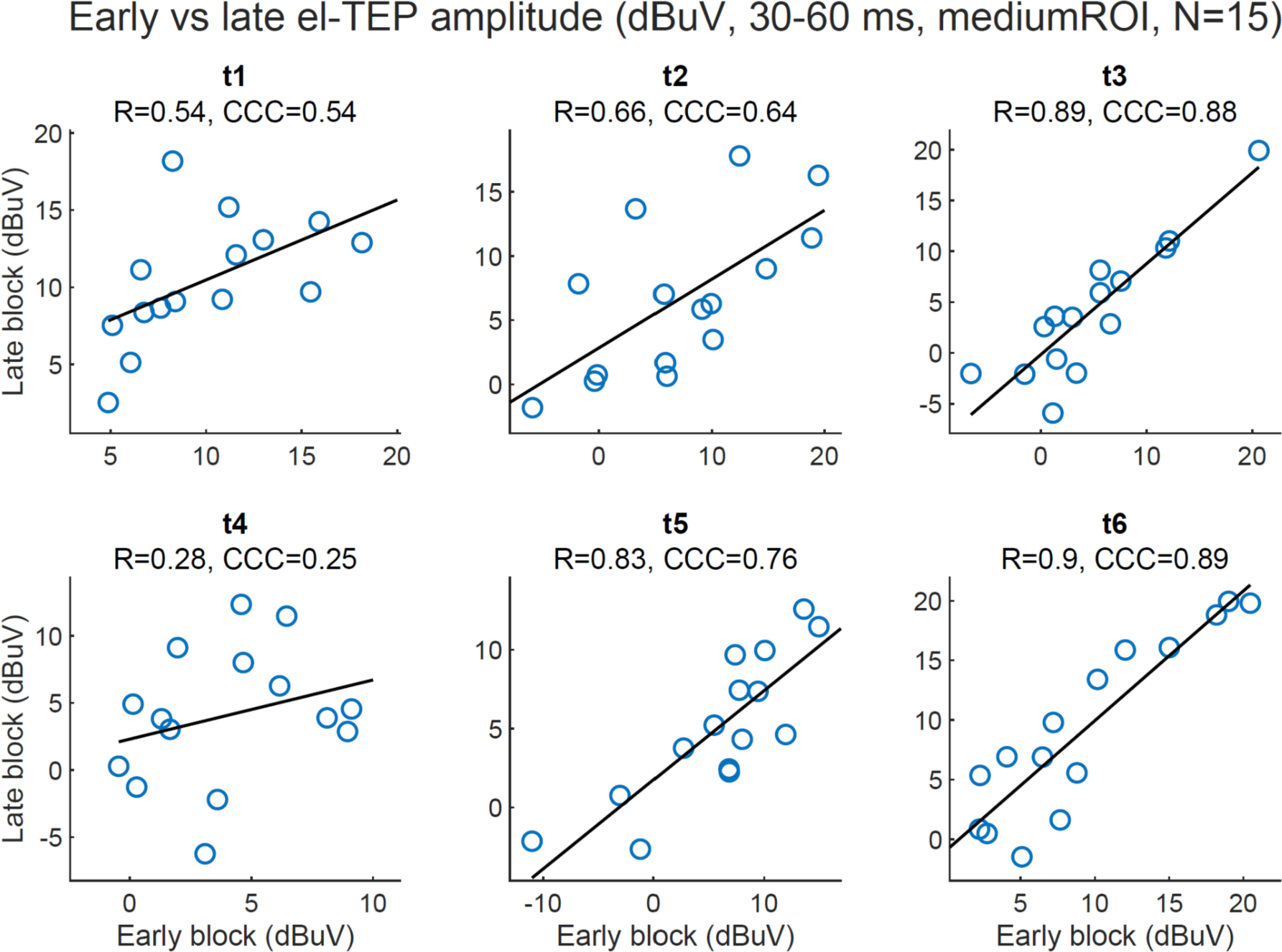
EL-TEP amplitudes (medium ROI, 30-60 ms time window, peak-to-peak) of the first and the second experimental block. Each dot represents one subject (N=15).

**Figure S11.**
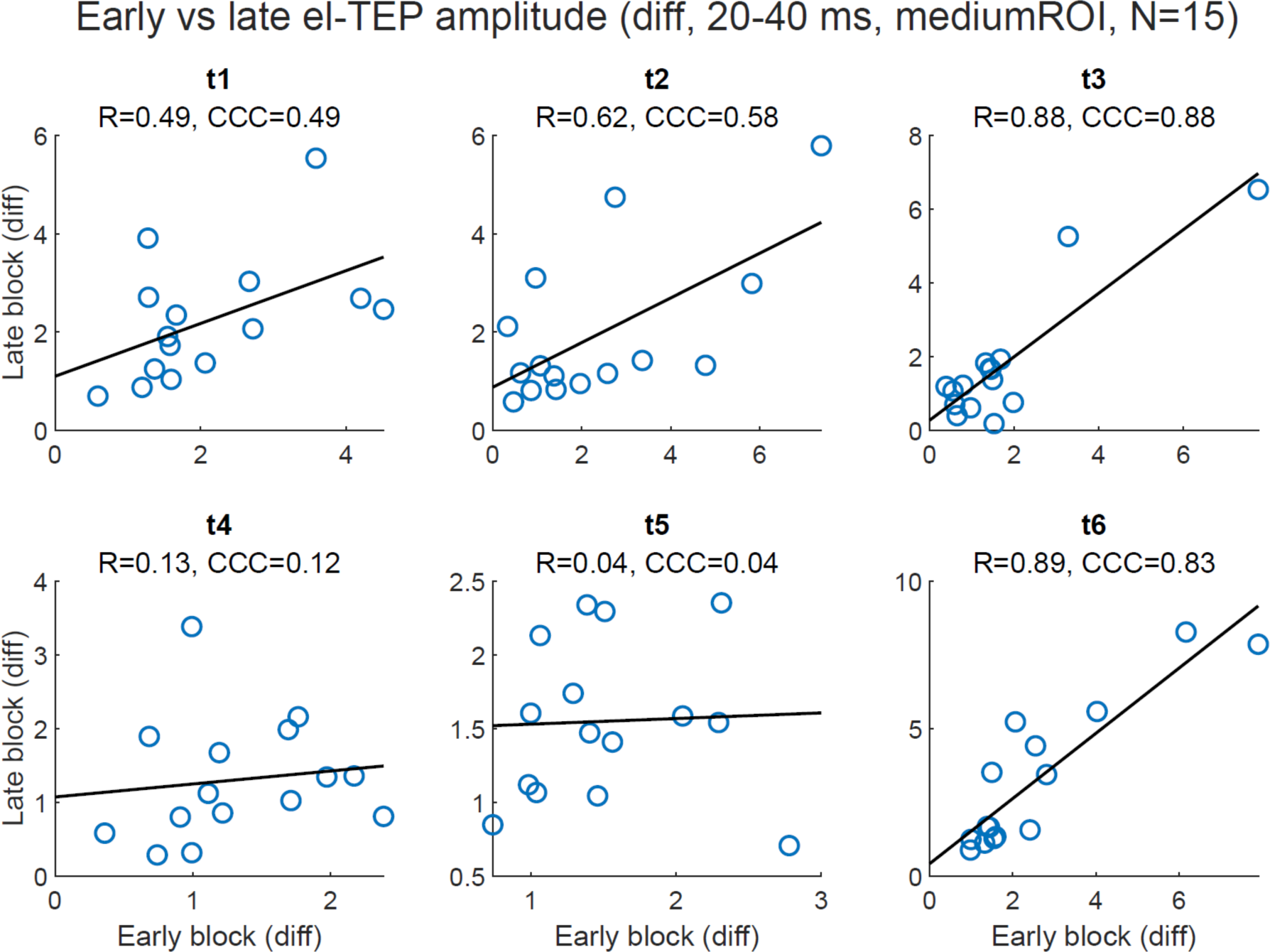
EL-TEP amplitudes (medium ROI, 30-40 ms time window, peak-to-peak) of the first and the second experimental block using linear (μV) peak-to-peak values (instead of dBμV). Each dot represents one subject (N=15).

**Figure S12.**
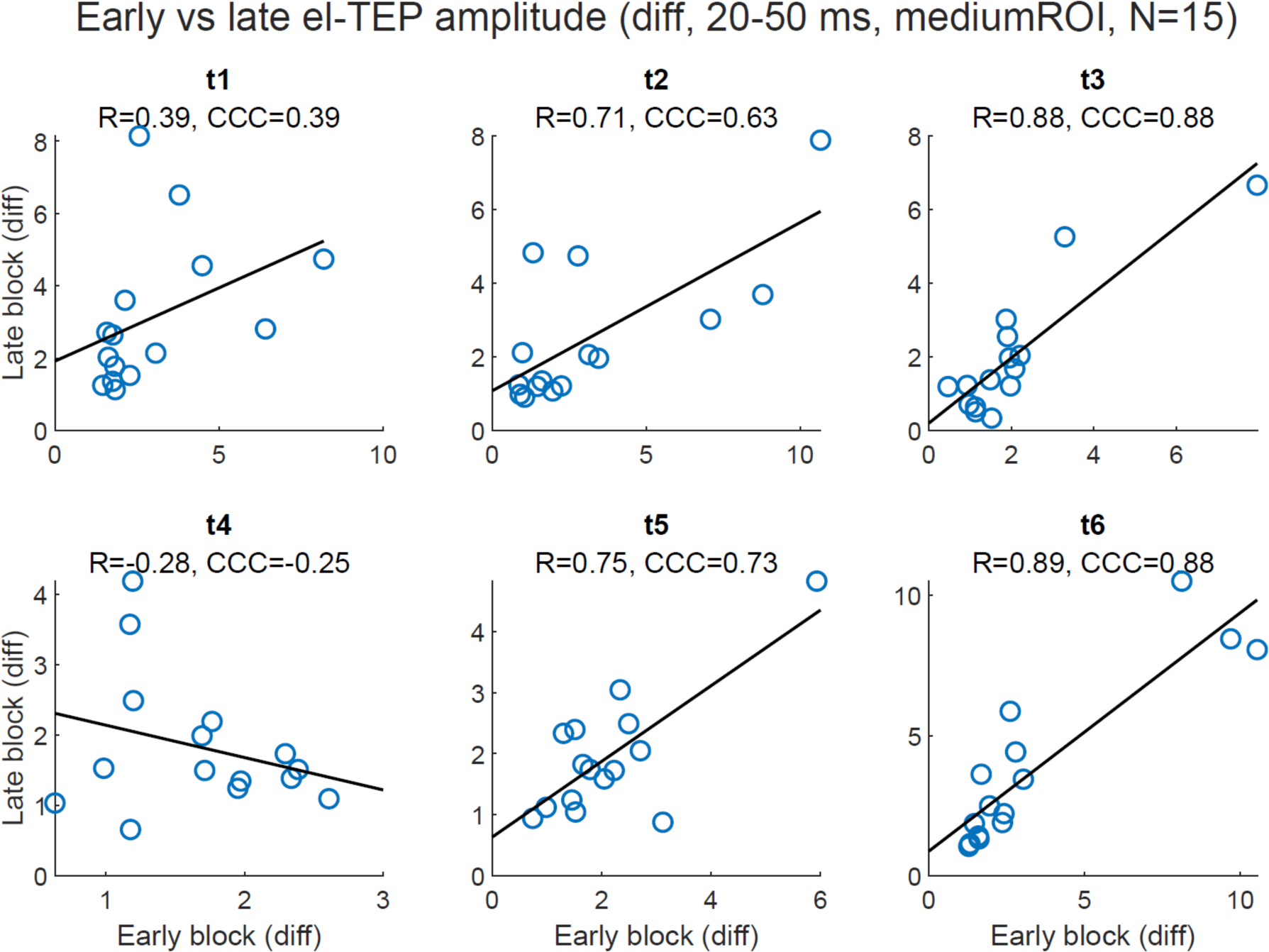
EL-TEP amplitudes (medium ROI, 20-50 ms time window, peak-to-peak) of the first and the second experimental block using linear (μV) peak-to-peak values (instead of dBμV). Each dot represents one subject (N=15).

**Figure S13.**
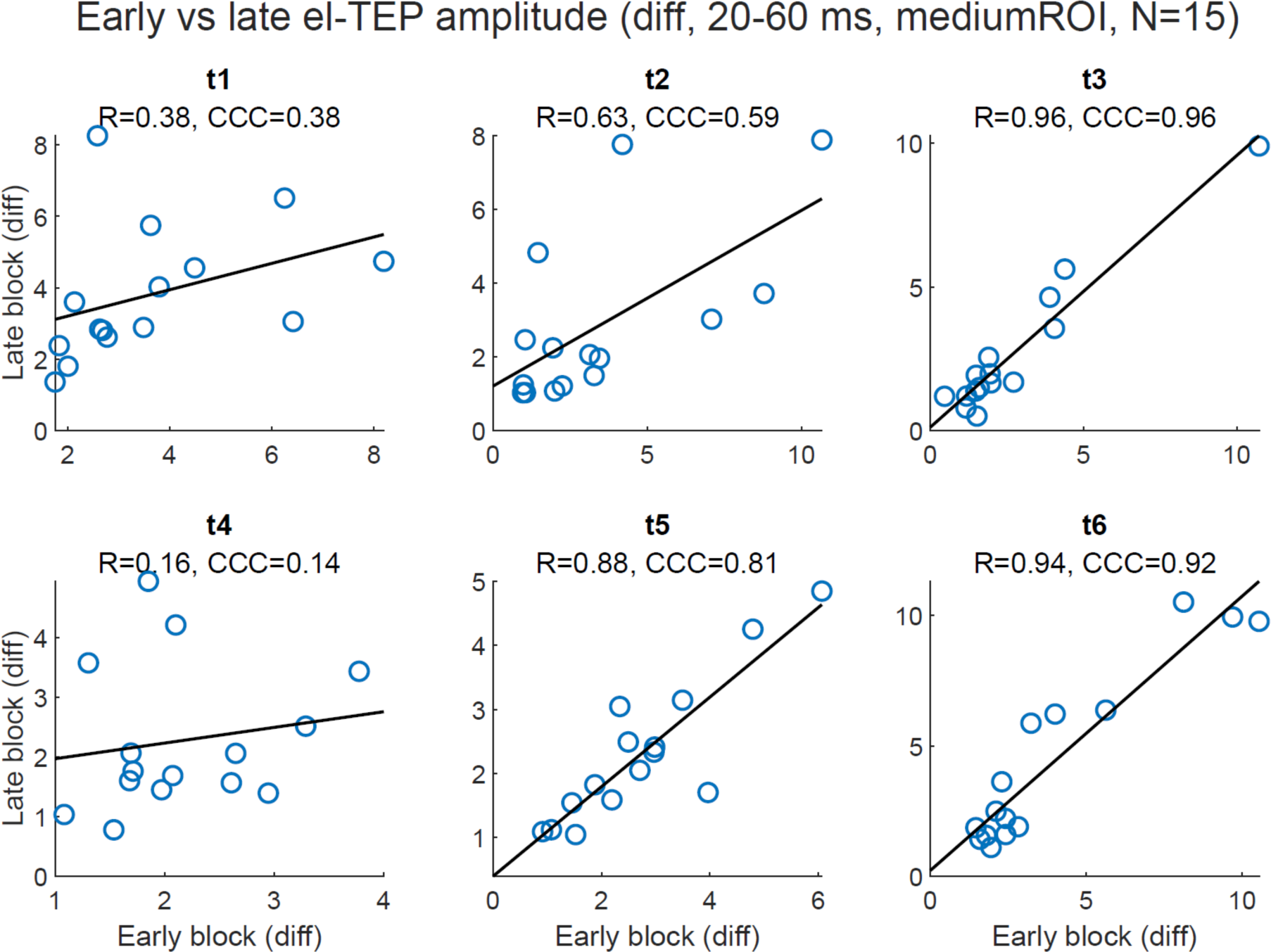
EL-TEP amplitudes (medium ROI, 20-60 ms time window, peak-to-peak) of the first and the second experimental block using linear (μV) peak-to-peak values (instead of dBμV). Each dot represents one subject (N=15).

**Figure S14.**
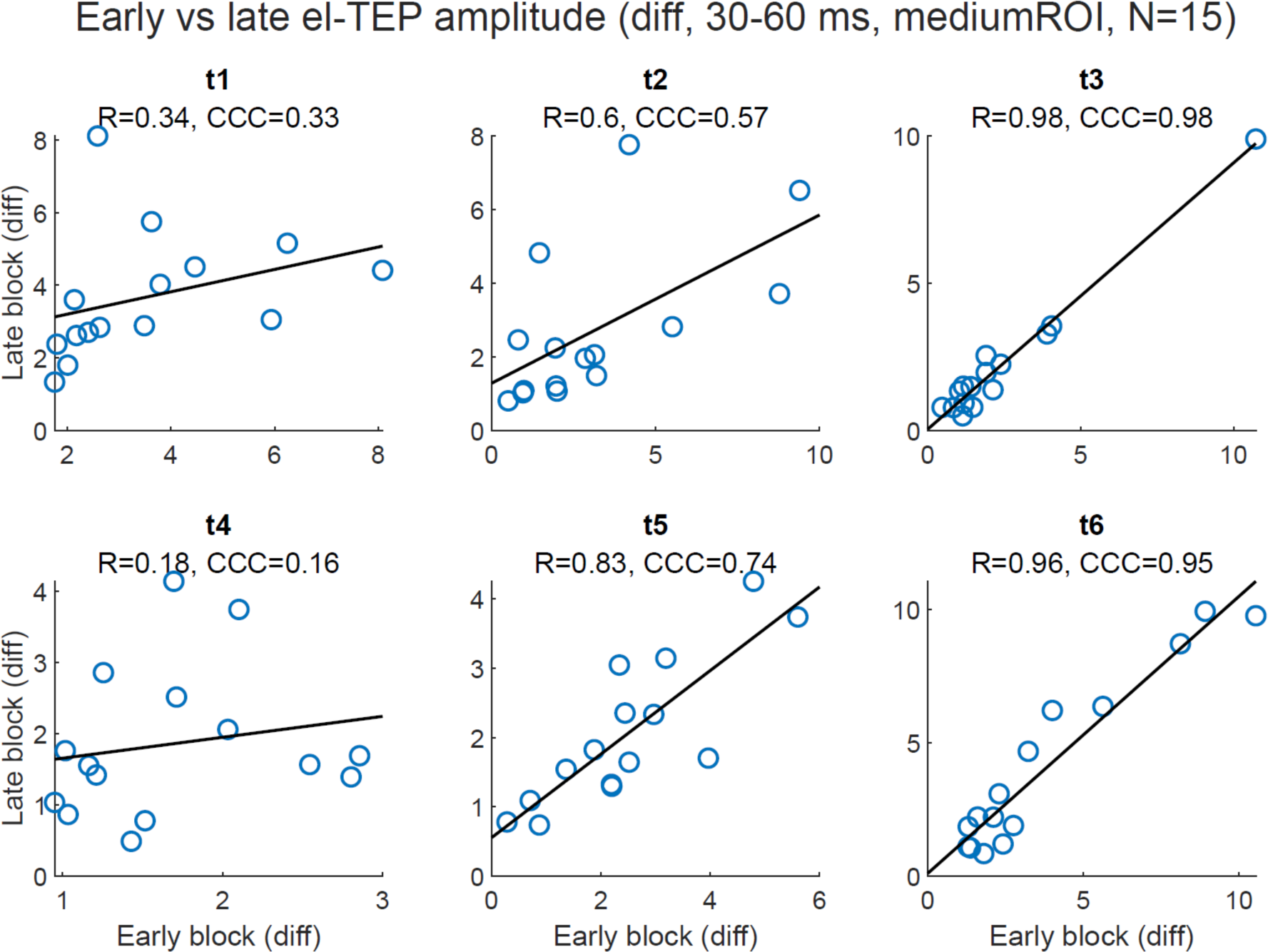
EL-TEP amplitudes (medium ROI, 30-60 ms time window, peak-to-peak) of the first and the second experimental block using linear (μV) peak-to-peak values (instead of dBμV). Each dot represents one subject (N=15).

**Fig S15.**
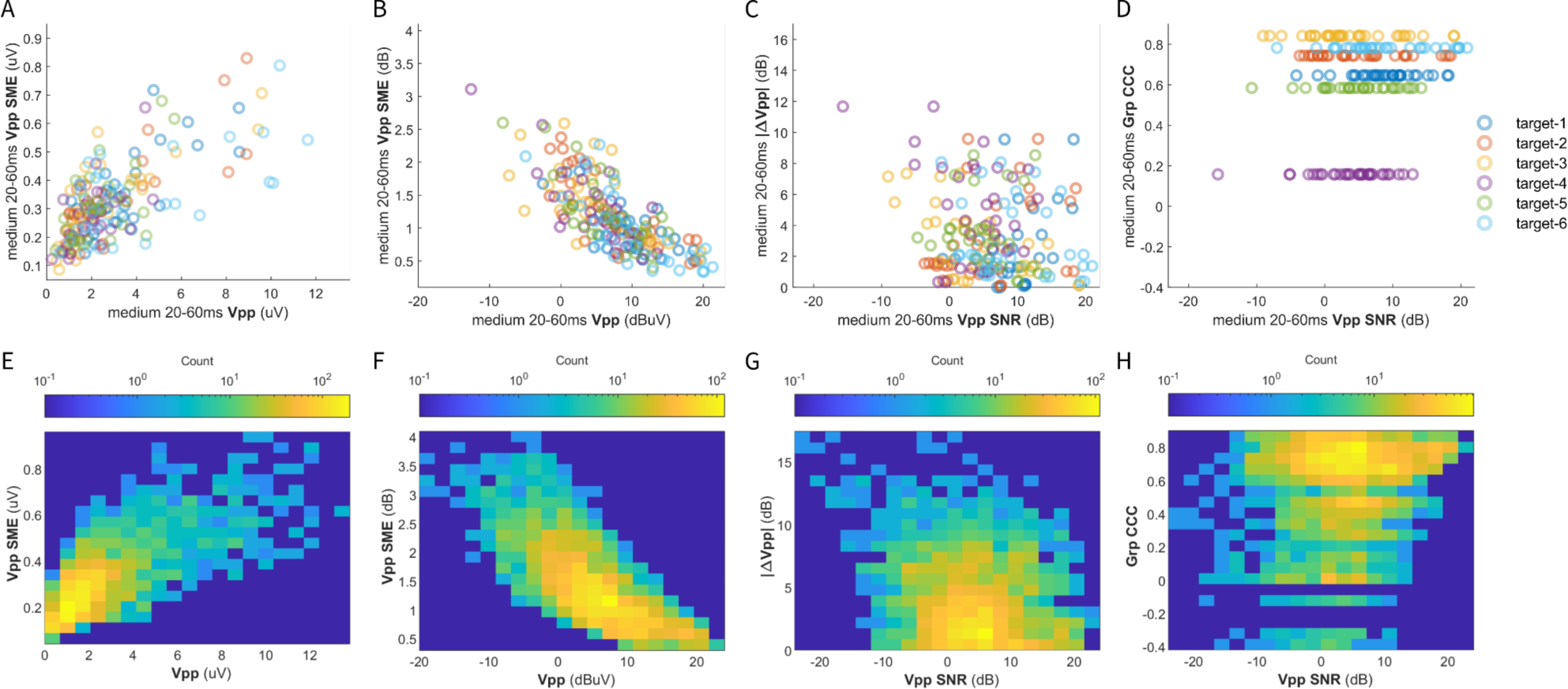
Standardized measurement error (SME) analysis results. A-D) scatter plots comparing within-block and across-block amplitude, variance, and reliability metrics for the medium-size sensor-space ROI, 20-60 ms time window, and full set of 150 trials within each block. A) scatter plot of within-block mean linear response amplitude (linear scale) vs within-block linear SME. B) scatter plot of within-block mean logarithmic response amplitude vs within-block logarithmic SME. C) scatter plot of within-block logarithmic SNR vs across-block response amplitude change. D) scatter plot within-block logarithmic SNR vs. across-block group-level CCC. E-H) histograms for the same variables as (A-D) but pooled across all full-trial-count sensor-space analytical variable combinations.

**Fig S16.**
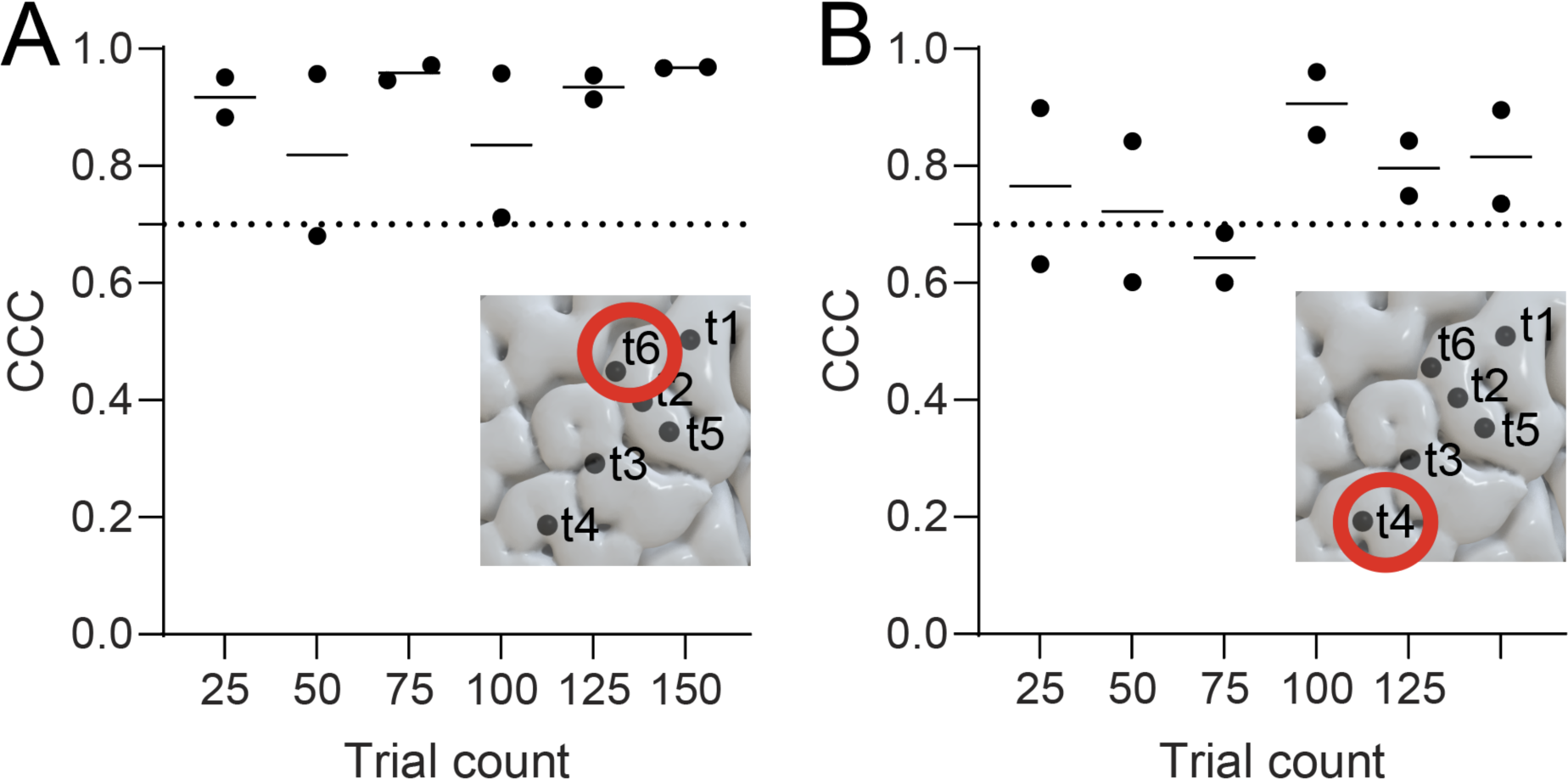
Impact of automated preprocessing on EL-TEP reliability. Automated preprocessing was run twice for identical trials and CCCs were computed using the two preprocessing results. The dots in each trial count condition represent the first and the second stimulation block. Automated preprocessing introduces some variability in CCCs, particularly noticeable in target t4.

**Fig S17.**
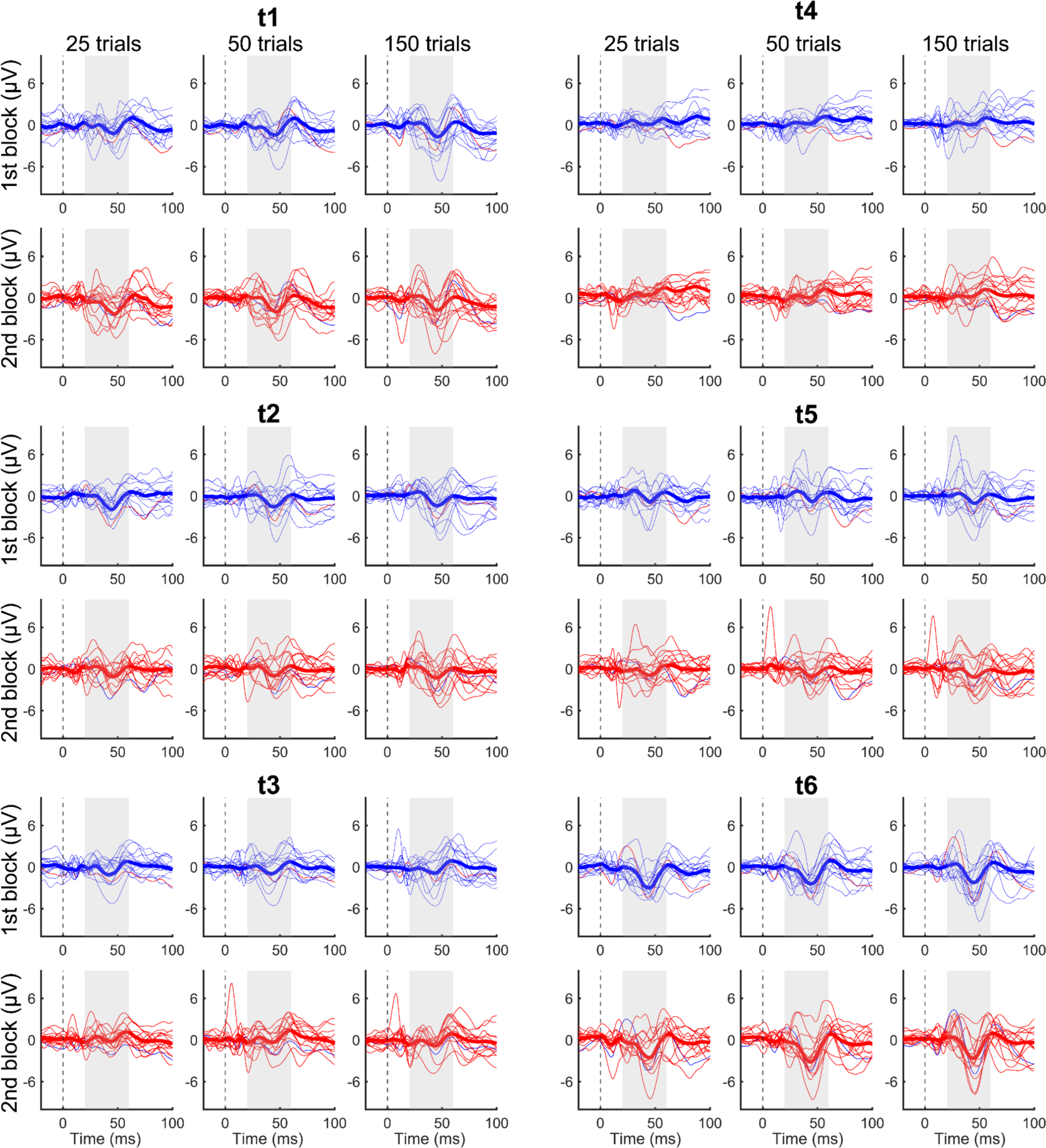
TEP traces across targets for trial count analysis. TEP traces for 25, 50 and 150 trials for the first and second stimulation block are shown for each stimulation target. Bold lines represent average TEP response across subjects.

